# Lab-in-the-loop therapeutic antibody design with deep learning

**DOI:** 10.1101/2025.02.19.639050

**Authors:** Nathan C. Frey, Isidro Hötzel, Samuel D. Stanton, Ryan Kelly, Robert G. Alberstein, Emily K. Makowski, Karolis Martinkus, Daniel Berenberg, Jack Bevers, Tyler Bryson, Pamela Chan, Yongmei Chen, Alicja Czubaty, Tamica D’Souza, Henri Dwyer, Anna Dziewulska, James W. Fairman, Allen Goodman, Jennifer Hofmann, Henry Isaacson, Aya Ismail, Samantha James, Taylor Joren, Simon Kelow, James R. Kiefer, Matthieu Kirchmeyer, Joseph Kleinhenz, James T. Koerber, Julien Lafrance-Vanasse, Andrew Leaver-Fay, Jae Hyeon Lee, Edith Lee, Donald Lee, Wei-Ching Liang, Joshua Yao-Yu Lin, Sidney Lisanza, Andreas Loukas, Jan Ludwiczak, Sai Pooja Mahajan, Omar Mahmood, Homa Mohammadi-Peyhani, Santrupti Nerli, Ji Won Park, Jaewoo Park, Stephen Ra, Sarah Robinson, Saeed Saremi, Franziska Seeger, Imee Sinha, Anna M. Sokol, Natasa Tagasovska, Hao To, Edward Wagstaff, Amy Wang, Andrew M. Watkins, Blair Wilson, Shuang Wu, Karina Zadorozhny, John Marioni, Aviv Regev, Yan Wu, Kyunghyun Cho, Richard Bonneau, Vladimir Gligorijević

**Author notes:** These authors contributed equally to this work (Nathan C. Frey, Isidro Hötzel, Samuel D. Stanton, Ryan Kelly, Robert G. Alberstein, Emily K. Makowski).

## Abstract

Therapeutic antibody design is a complex multi-property optimization problem with substantial promise for improvement with the application of machine-learning methods. Towards realizing that promise, we introduce “Lab-in-the-loop,” a new approach that orchestrates state-of-the-art repertoire mining methods, generative machine learning models, multi-task property predictors, active learning ranking and selection, and *in vitro* experimentation in a semi-autonomous, iterative optimization loop. By automating the design of antibody variants, property prediction, ranking and selection of designs to assay in the lab, and ingestion of *in vitro* data, we enable an end-to-end approach to developing computationally-informed therapeutic antibody design pipelines. We apply lab-in-the-loop to eleven seed antibodies obtained via animal immunization with four clinically relevant antigen targets: EGFR, IL-6, HER2, and OSM. Over 1,800 unique antibody variants are tested throughout four rounds of iterative optimization identifying 3–100× better binding variants for all targets and 10/11 seeds, with the best binders exceeding 100 pM affinity, demonstrating a process by which end-to-end machine learning can be developed for therapeutic antibody development.

## 1 Main

Discovery of therapeutics is a fundamental driver of the extension of human life and health-span. Antibodies represent one of the most versatile classes of therapeutic modalities. The genetic modularity and biological mechanisms for rapid diversification underlying their production and maturation enable the jawed vertebrate immunological response to recognize almost any target molecule of interest, including proteins, peptides, and small molecules [1, 2]. As therapeutics, antibodies can be leveraged for signaling pathway modulation, immune system engagement, or to deliver drug payloads with high specificity, creating strong demand for new antibodies, leading to 58 new FDA-approved antibody therapeutics between 2019–2023 [3]. However, antibody discovery and engineering remains a challenging process, adding to high development costs that influence the final cost of these medicines.

Therapeutic antibody discovery campaigns commonly begin with the immunization of animals with the target antigen, eliciting an immune response that can be quantified by sequencing and analysis of the B cell repertoire before and after immunization [4]. In addition to repertoire sequencing, the immune response can be processed via hybridoma generation or single B cell screening yielding initial antibody hits, which we refer to as *seeds*. Alternatively, large libraries of 10^9^–10^11^naive antibody sequences can be expressed via display technologies (e.g., yeast or phage) and panned against the antigen directly *in vitro* to yield initial antibody seeds [5]. These initial seeds then must be optimized for antigen binding affinity, functional activity, immunogenicity, pharmacokinetics, expression, stability, solubility, viscosity, and aggregation propensity to satisfy the many criteria required for an antibody to be a viable therapeutic molecule [6, 7]. For therapeutic antibodies, a common optimization task is affinity maturation, where carefully chosen mutations are incorporated to increase the affinity of a molecule without introducing liabilities. These engineering efforts are primarily aimed at increasing the biological efficacy of the molecule. Success in affinity maturation efforts can be considered through both the fold increase in affinity and the binary success of achieving a specified threshold of affinity maturation. We consider ≥3× increases in affinity as our success criteria as this increase exceeds instrument noise by an order of magnitude and represents an increase at which the biological efficacy of increasing affinity can be determined. The high-dimensional search space for antibody sequences has largely precluded first-principles approaches to therapeutic antibody design, including affinity maturation, and the large cost, long timelines, and linearity of existing processes are impediments for the rapid development of new therapeutics [8] as they cannot effectively manage the balance of multiple properties. Machine learning (ML) and Bayesian Optimization (BayesOpt) are natural approaches for navigating the sparsely populated, high-dimensional space of functional protein sequences [9–15]; indeed, language models trained on protein sequence data have been used to guide directed evolution to affinity mature antibodies [16–19]. However, this sparsity renders the *de novo* discovery of initial sequences for optimization exceptionally challenging. Thus, the facile identification of diverse antibody seeds via animal immunization (or *in vitro* libraries) and powerful optimization capabilities of ML and BayesOpt algorithms exhibit reciprocal strengths spanning the entire drug discovery process. Crucially, for drug discovery campaigns, ML methods must respect unique constraints for therapeutic molecules (e.g., developability, specificity, potency, expression, efficacy) and augment state-of-the-art molecular engineering approaches [6]. Prior work in ML for antibody design shows the immense promise of these approaches [16, 20]. However, in our hands these methods result in antibody variants that do not improve binding affinity, do not diversify the lead candidates, or introduce germline-reverting mutations that abrogate affinity. Unlike prior work, our optimization campaigns begin with reasonably optimized and viable clones from state-of-the-art *in vivo* discovery campaigns. Because of the stringent requirements and the complexity of the multi-property optimization problem outlined above, *de novo* protein discovery methods [21–23] do not obviate the need for optimization. Our approach takes steps towards respecting important therapeutic constraints [6], demonstrates generality across antigen targets and epitopes, and enables semi-autonomous antibody engineering through a combination of state-of-the-art experimental and computational methods. Here, to address the limitations of both traditional discovery/engineering and prior ML-based design of antibodies, we introduce the “Lab-in-the-loop” (LitL) system, which combines the complementary advantages of both to systematically advance nonlinear sequence optimization.

We report results for four therapeutically relevant and biologically interesting antigen targets: Epidermal growth factor receptor (EGFR) [24], Interleukin-6 (IL-6) [25], Human epidermal growth factor receptor 2 (HER2; ERBB2) [26], and Oncostatin M (OSM) [27]. By leveraging a semi-automated ensemble of generative models coordinated by a property prediction model for ranking and selection of designs, we achieve improvements across affinity, expression, and developability. We discover between 3–100× better binding molecules for all four targets and for ten out of eleven seeds, starting with antibodies from immunization discovery campaigns and select initialization data, primarily from repertoire mining efforts. We rationalize our results via experimental structure determination of 8 seed candidates and designs, as well as biophysics-based computational modeling of binding design complexes, showing that our designs preserve important interactions responsible for binding, while introducing up to eight new mutations that improve binding affinity. Our results demonstrate the powerful generalization capabilities of LitL to harmonize active learning approaches including ML and BayesOpt with at-scale antibody characterization, towards the goal of producing a system for the computationally-informed design of therapeutic antibodies.

## 2 Results

### 2.1 Lab-in-the-loop is a general machine learning system for autonomous drug discovery

LitL is a new approach that orchestrates loosely coupled ensembles of generative models, property prediction oracles, a ranking and selection algorithm, and experimental assays, all encompassed in an active learning loop to achieve iterative optimization of antibody lead candidates (Fig 1**a**). Unlike prior work on biomolecular design with ML, which does not reflect the constraints and rigorous standards for success needed in drug discovery campaigns [6, 28, 29], LitL explicitly incorporates state-of-the-art experimental methods, including animal immunization and repertoire mining, optimizes from viable lead candidate molecules, and operates across the human proteome of druggable antigen targets. In this paper, we focus on therapeutic antibody sequence space, although LitL is extensible to other therapeutic modalities. The selected example targets of EGFR, IL6, HER2, and OSM are of great interest in the treatment of cancer, inflammation, and autoimmune diseases [24–27]. Our active learning process is comprised of four rounds, over which we produce over 1,800 antibody designs (Table 1).

**Table 1:** Lab-in-the-loop statistics.

| Design round | Total designs | Expressors ($> 0.01$ mg) | Binders ($> 4pK_D$) | Better binders ($> 3\times$) |
| --- | --- | --- | --- | --- |
| 1 | 533 | 412 | 177 | 12 |
| 2 | 520 | 499 | 420 | 24 |
| 3 | 771 | 681 | 508 | 135 |
| 4 | 305 | 291 | 258 | 65 |

**Fig. 1:**
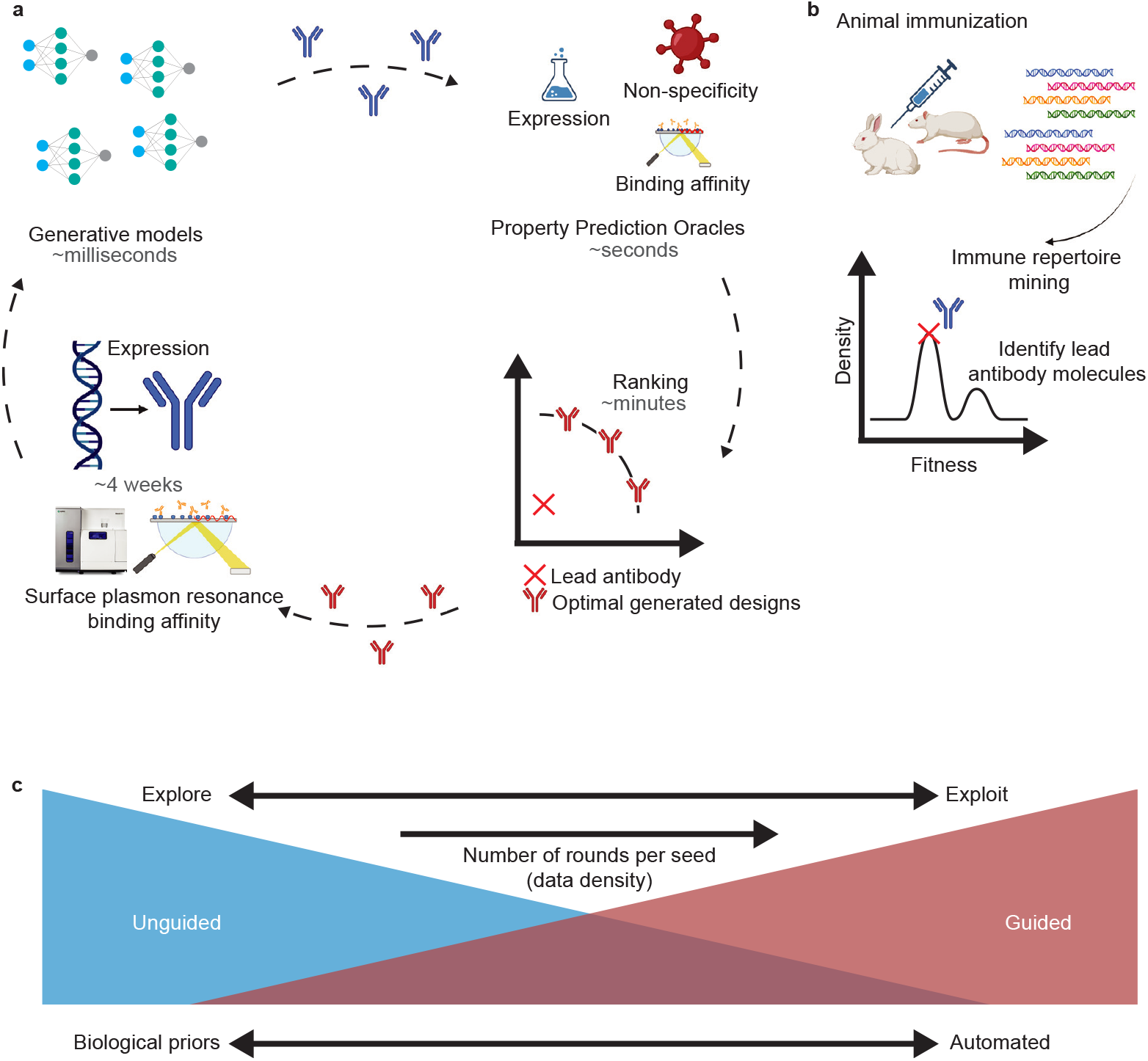
Lab-in-the-loop autonomous therapeutic antibody optimization. **a**, Generative models produce a diverse *in silico* library of antibody sequence designs. Designs are ranked by property prediction oracles that are chosen, tournament style, based on their leaderboard performance evaluated on a hold-out split from the previous round of experiments. Predicted properties are used to rank designs to maximize the expected improvement compared to the lead antibody. Designs are expressed and binding affinity is measured to antigen targets with surface plasmon resonance (SPR). Lab data is used to re-train generative models and property prediction oracles for active learning. **b**, Lead antibody molecules are the best identified starting points from *in vivo* animal immunization and immune repertoire mining, although our approach can be initialized from any antibody discovery method. **c**, Ensembling unguided and guided generative models allows the LitL framework to dynamically explore and exploit regions of antibody sequence space. When data is sparse, LitL performs unguided exploration of nearby sequence space constrained by biological priors (e.g., conserve framework regions for humanness and immunogenicity, mutate CDRs for affinity modulation). As active learning proceeds, labeled data is collected, and models are retrained, the molecule design process moves towards guided generation and “exploitation” of Pareto-optimal sequence space.

#### 2.1.1 Seed antibodies are obtained from *in vivo* discovery campaigns

Ten of our eleven seed antibodies are obtained from state-of-the-art *in vivo* immunization of genetically engineered animals and immune repertoire mining techniques (Fig. 1**b**) [4]. Repertoire mining produces natural, developable seed molecules and additionally provides unlabeled local neighborhoods (in sequence space) of likely binders around the seeds. Though we find repertoire mining data invaluable for initial training sets of generative and property prediction models, the LitL approach is fully general and starting seeds can also be obtained from *in vitro* discovery campaigns [30]. To demonstrate the versatility of our methods and enable comparative benchmarking, we include hu4D5 (trastuzumab) as an eleventh seed.

Starting from a seed molecule that we aim to improve, the loop begins with the selection of a limited number of additional sequences from the local sequence space surrounding the seed. These sequences are primarily from repertoire mining and initialize our LitL machine-learning models, enabling the creation of a diverse library of sequences (i.e., the variants of the seed sequence) that is rich in high-quality candidate molecules. Further details on seed selection and repertoire mining are presented in Section 4.4.2 of the Methods.

#### 2.1.2 Generative models produce libraries of candidate molecules

Generative models are trained on protein sequence data with the aim of emitting novel, diverse, functional proteins that are improved relative to sequences from *in vitro* or *in vivo* discovery and design campaigns. While *in vitro* and *in vivo* methods are restricted to sequences that are present in the display library or animal immune repertoire (which may require significant engineering before they are suitable for clinical trials), ML methods can generate near-limitless sequence diversity, optimize sequences with respect to multiple desirable properties, and enforce constraints on chemical liabilities and other molecular assessment criteria that would otherwise prevent molecules from advancing to clinical trials.

Generative models have various trade offs, generally between sample quality and sample diversity. To address the limitations of any particular generative model, we developed purpose-built methods for unguided sampling, sequence diversification, and hit expansion (e.g., Walk-Jump Sampling [31]) and separate generative methods for guided multi-property optimization (e.g., LaMBO-2 [32], DyAb [33], and Property Enhancer [34]). We include design proposals from multiple methods in each round, with the purpose of diversifying our design pool and maximizing the Noisy Expected Hypervolume Improvement (NEHVI) *acquisition function* [35], as each method explores different parts of the antibody sequence space (see Supplementary Figures B3, B4, B5, B6).

We adaptively allocate experimental capacity to each method by combining the output of each method into a single combined library that is subsequently filtered and ranked by our active learning framework (i.e., global selection). In each round, all generative methods may produce up to 30,000 designs per lead molecule. This constraint is imposed to prevent any one method from dominating global selection by brute force, promote sequence diversity by attenuating the sampling biases of individual models, and to limit the computational cost of downstream *in silico* quality control. Details of these generative methods, their architectures, and training datasets, are presented in Section 4.1.1 of the Methods and their respective references.

#### 2.1.3 Diverse molecule libraries are filtered and ranked for active learning selection

Given a high-quality library of designs, we then estimate their therapeutic properties to determine which sequences to synthesize. In particular, generated designs are annotated with predicted properties encompassing expression, binding affinity, and non-specificity. All properties have associated binary labels (e.g., 1:1 binding to target antigen is or is not detected by SPR at a fixed concentration) modeled by binary classifiers, and expression yield and binding affinity are modeled as scalars by regression models. Out-of-distribution (OOD) detection methods introduced in [31] and additional quality control filters (e.g., chemical liability motifs) are also used to assist the discriminator by excluding very anomalous sequences or those with easily identified developability issues. Further details of these oracles, OOD detection methods, and quality control filters are presented in Sections 4.1.3, 4.1.4, and 4.1.5 of the Methods and their respective references.

The predicted properties for each design that passes quality control filters are then used to produce a global ranking of the designs that are predicted to perform best in laboratory experiments. The ranking model is the global arbiter of design selection. This systems-level design allows LitL to incorporate many diverse ideas and methods, while ensuring that all experimental successes contribute to improving the ranking model and oracles, which in turn improves the generative models, creating a flywheel of continuous improvement. The ranking algorithm picks designs that are *non-dominated*, that is, designs on the Pareto frontier of property space. We want to avoid expensive experimental assays for designs that are strictly dominated, i.e., designs for which we already have labeled antibody candidates with properties at least as good or better in all respects than the candidate design.

We select designs using the NEHVI acquisition function [35] to account for the noise in property measurements, promote diversity in property space, and balance the explore-exploit tradeoff induced by the conflicting objectives of improving the generalization capabilities of the discriminative models while still producing high-quality candidate molecules as quickly as possible. Further details on the acquisition function and ranking are given in Section 4.1.6 of the Methods.

#### 2.1.4 Experimental workflows drive the active learning loop

Given the complex nonlinear mapping between protein sequence and properties, the challenge of acquiring experimental measurements at scale, and the critical importance of high quality data for training and evaluating models, the cornerstone of LitL is an efficient experimental pipeline capable of producing hundreds of high-resolution data points in parallelized 4-6 week timelines. To enable the accelerated timelines of LitL, we developed an optimized antibody production pipeline utilizing Gibson-assembled linear fragments (GLFs), as previously described [36]. Briefly, variable region sequences are synthesized as linear DNA fragments (Twist Bioscience) and subsequently integrated into GLFs containing all elements required for transient expression at the 1mL scale utilizing HEK293 cells. The linear DNA based expression workflow trims multiple cloning steps, saving significant time and cost while also being highly amenable to automation. Binding affinity of the designed and lead antibodies is measured by surface plasmon resonance (SPR). Further details on experimental methods are presented in Section 4.4 of the Methods.

#### 2.1.5 Active learning

LitL proceeds as an active learning loop over multiple design rounds described above. In the first round, minimal labeled data is available. In each successive round, we perform multi-objective Bayesian optimization: labeled data is collected and used to retrain the generative models and the predictive multi-task model, which is then used to rank designs and decide what data to collect next. This online procedure progressively improves both the models and the quality of proposed designs. We report four rounds in this work. Further details on the ranking and selection procedure are presented in Methods Section 4.1.6.

### 2.2 Lead candidates are affinity matured via iterative optimization

To demonstrate the utility of LitL, we applied our process to the optimization of eleven antibodies against four clinically relevant targets. For simplicity, our top line metric of success for a design is a strict threshold of at least 3× improvement in measured binding affinity compared to the lead antibody. We represent binding affinity as a negative log transform of the equilibrium dissociation constant,

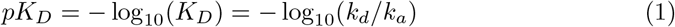

where *k*_*d*_ is the dissociation rate (off rate) and *k*_*a*_ is the association rate (on rate). Improved affinities correspond with higher *pK*_*D*_, with a greater than 3× affinity improvement equivalent to Δ*pK*_*D*_ *>* 0.477. In the first round of LitL, we initialize our computational methods with repertoire-based designs and limited computationally-informed designs, identifying 3× binders for 3/11 seeds (2% of total designs; Fig. 2a). By the fourth round, LitL produces a design library of *>*21% 3× better binders (Fig. 2a). Figure B7 shows the cumulative number of labeled designs for each seed over design rounds.

**Fig. 2:**
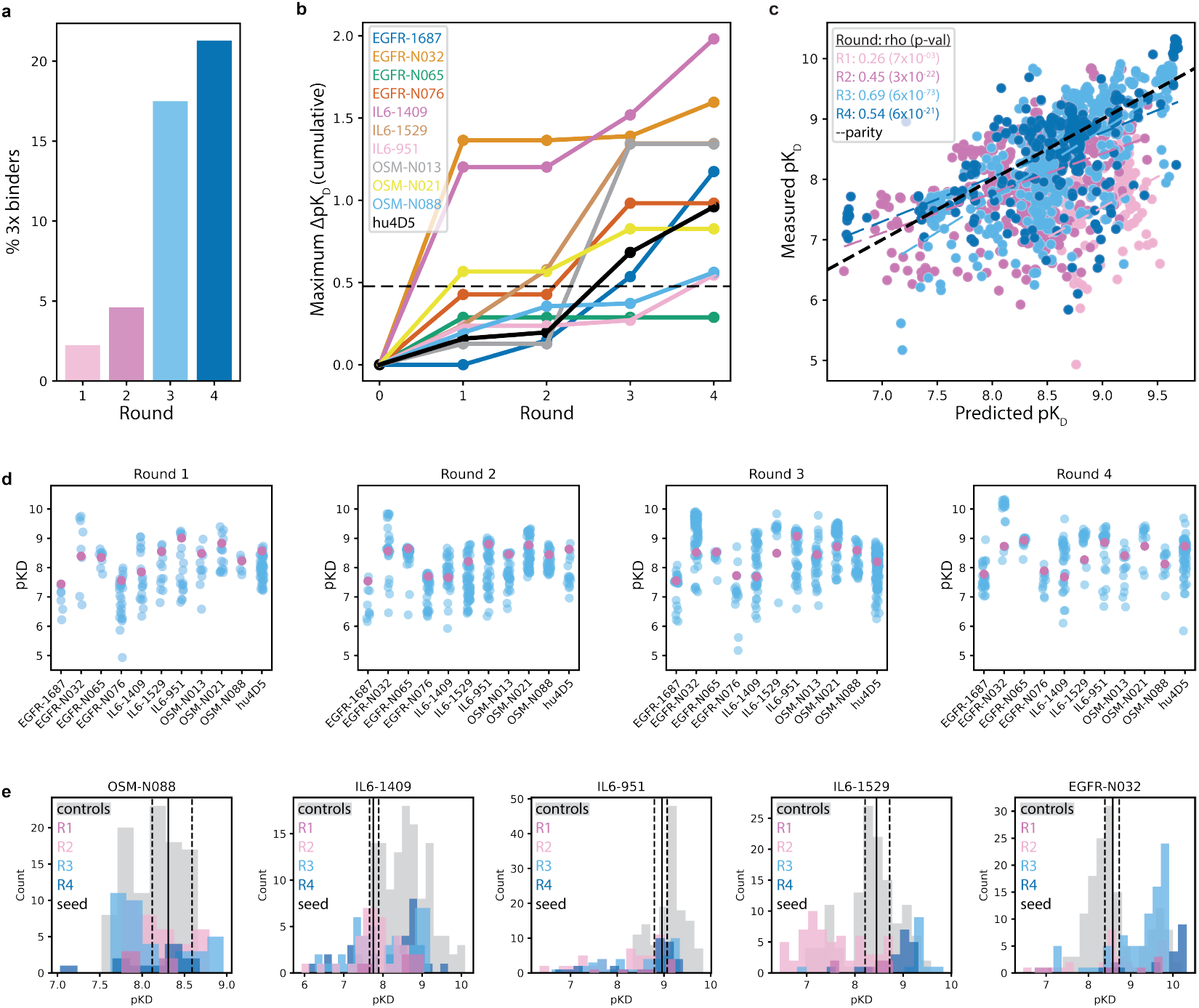
Therapeutic antibody lead optimization via affinity maturation with lab-in-the-loop. **a**, Lab-in-the-loop achieves compounding improvements in rate of generating 3×-tighter binding variants over successive rounds of design. **b**, Cumulative maximum improvement in binding affinity over four rounds for eleven different seed antibodies across four different antigen targets. The dashed horizontal line indicates a 3× improvement in binding affinity. **c**, Predicted versus measured *pK*_*D*_ for all designs. **d** Absolute measured *pK*_*D*_ (higher is better) of designs (blue) and lead molecules (pink) over design rounds. **e** Absolute measured *pK*_*D*_ for five seeds and designs over four rounds compared to baseline control measurements (Section 2.3). The solid vertical line represents the average *pK*_*D*_ measurement of the seed while the dashed vertical lines represent the minimum and maximum *pK*_*D*_ measured for each seed throughout all Lab-in-the-loop rounds.

Evaluating the highest affinity clone from each lead, LitL successfully discovers a 3× better binder in ten out of eleven cases (Fig. 2b). For five of the seeds, LitL yielded better binders with Δ*pK*_*D*_ *>* 1, which denotes a 10× increase in binding affinity. In Figure 2c we show predicted versus measured *pK*_*D*_ for all designs over the four rounds, revealing improved correlations with progress through the rounds. Figure 2d shows the absolute measured *pK*_*D*_ for designs (in blue) over rounds and their respective seeds (in pink). See Figures B8 and B9 for alternative visualizations.

### 2.3 Comparison of LitL outputs to generative baselines

To benchmark the performance of our ML models, we compared LitL designs against a randomized baseline “control” informed by state-of-the-art *in vivo* immunization and *in silico* repertoire mining techniques (we separately discuss other generative protein design methods in Section A). Control designs were generated by first identifying Round 1 repertoire picks for five seed antibodies possessing comparable or improved affinities, then randomly sampling from these sets of acquired mutations to generate novel seed variants, taking care to match both the total number of antibodies evaluated and the distribution of mutational loading to the aggregate of designs generated by ML methods in Rounds 2–4.

Affinity measurements for the control samples had comparable distributions to our machine-learning-informed designs (Figure 2e). For three of five seed antibodies, the highest affinity improvement was achieved via ML methods in either the third or fourth round of optimization. For two out of five seed antibodies, the highest affinity improvement was achieved in the control antibodies. We note, however, that control antibodies were generated without regard for developability constraints or expression yield optimization, in contrast with model designs which were generated with constrained multi-property optimization.

### 2.4 Multi-property optimization enables therapeutic antibody design

In addition to initializing our optimization efforts from developable, natural antibodies derived from *in vivo* immune repertoire mining, we further constrain our designs to account for non-specificity, expression yield (which impacts downstream characterization assays that require specific amounts of protein material), and *in silico* developability risks. This differentiates our efforts from previous work [15] that similarly applied iterative rounds of affinity/expression co-optimization, but initialized from naive (pre-”seed”) antibodies [16] and proceeded without consideration of these additional constraints. In Fig 3**a** we show the lead candidate molecules in dark blue and the best binding variant designs for each seed in white, after computing *in silico* metrics with the *Therapeutic Antibody Profiler* (TAP) [37]. These metrics, derived from the variable domain antibody sequence, flag sequences that are out-of-distribution with respect to the *Therapeutic Structural Antibody Database* (Thera-SAbDab). All but one of our best binding designs are within the TAP guideline ranges; one design is just over the threshold for positive and negative charge patches (PPC/PNC) in the CDRs. LitL avoids drastic biochemical and biophysical changes that would induce avoidable developability risks, while achieving many-fold improvements to binding affinity.

**Fig. 3:**
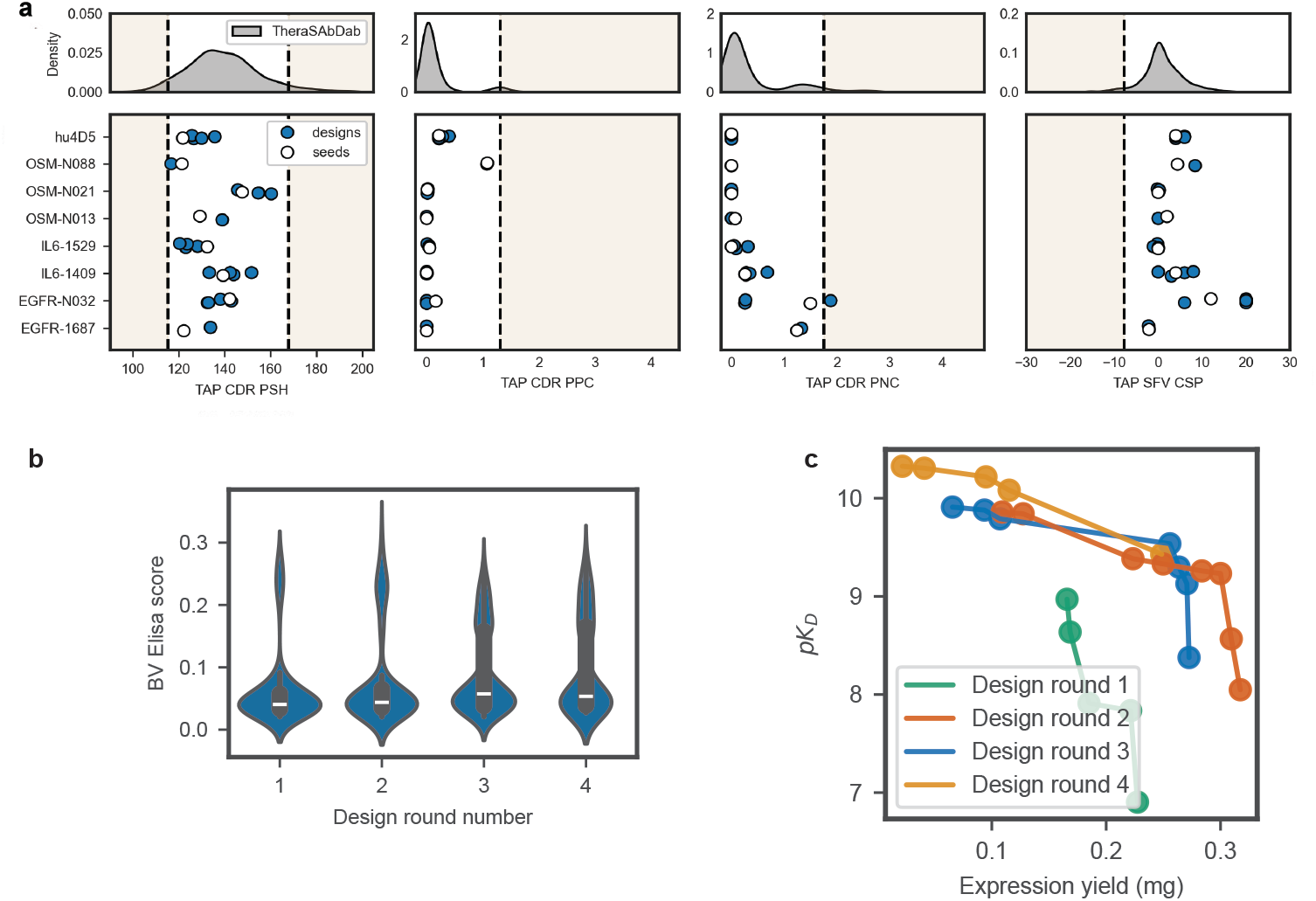
Multi-property optimization of lead candidates includes improving expression yield and *in silico* developability properties. **a**, All but one of the tightest binding design variants stay in acceptable ranges for developability guidelines given by the Therapeutic Antibody Profiler. Abbreviations: PSH - Patches of Surface Hydrophobicity; PPC - Patches of Positive Charge; PNC - Patches of Negative Charge; SFV CSP - Structural Fv Charge Symmetry Parameter. **b** Predicted BV ELISA scores for all designs, from a surrogate model trained on BV ELISA data, a measure of non-specificity. BVE scores *<* 1.0 indicate no non-specificity risk. As viable therapeutic starting points, all seeds have experimentally measured values within this range. **c**, Pareto frontier of non-dominated designs on the *pK*_*D*_ versus expression yield property landscape for each design round. See Supplementary Figure B13 for all data points.

To estimate risk of non-specific binding interactions of our designs, we developed a surrogate model trained on internal BV ELISA data. BV ELISA [38] is an assay developed to assess non-specific binding (further details are provided in Section 4.4). A BV ELISA score *>* 1.0 indicates a non-specificity risk; we confirm that all of our designs fall below the predicted risk threshold (Fig 3**b**). For 67 designs, we also engineer out chemical liabilities while maintaining or improving binding affinity (Fig B12).

Chemical liabilities include deamidation (‘NG’) and glycosylation (‘N*[ST]’) motifs, and other easily recognizable substrings that pose developability risks. The per-round Pareto frontier of designs with respect to expression concentration and binding affinity (Fig 3**d**) illustrates how successive design rounds expand the Pareto frontier, finding non-dominated designs. Though in this work we primarily illustrate co-optimization of two properties (filtering, not optimizing, for others), the ranking and selection approach of LitL can accommodate arbitrary numbers of properties simultaneously, with efficacy limited only by the accuracy of predictive models and the ability to acquire sufficient quantities of experimental data to train and evaluate them.

### 2.5 Modeled interface structures of designed antibodies demonstrate epistatic interactions

As all designs reported in this work were produced with sequence-only models, we considered whether their effects on affinity could be rationalized structurally (Fig 4). We first investigated two designs that achieved *>*4× improved binding (Δ*pK*_*D*_ = 0.688 (ERBB2-P00873, 5 edits, 1 insertion) and 0.638 (ERBB2-P00965, 8 edits, 1 deletion)) relative to the anti-HER2 antibody trastuzumab (hu4D5). From 10 independent minimizations of the native crystal structure (PDB ID: 1N8Z), we determined a union set of 17 antigen-contacting residues in the CDR loops through energetic analysis with Rosetta [39, 40] (Fig 4**a**). Modeling the singular edit at an existing heavy-chain contact (HC:N65Q in CDRH2)—a region with highly dynamic interactions (alternative rotamers in 4/10 simulations for HC:Y67, 3/10 for Ag:E558)—revealed that HC:N65Q loses all direct contact with HER2, but instead stabilizes the major rotamer of HC:Y67 via a new hydrogen bond, eliminating previous steric impact to the sidechain conformations of HC:R69 that in turn influences HC:Y40, resulting in a collective region-wide stabilization of interfacial contacts clearly visible in 4**a** (inset), demonstrating that direct optimization of interface positions indeed occurs and may exhibit epistatic effects on binding [41]. However, all other edits made across both designs (Supplementary Figure B14) are distinct from the native contacts, with many outside the CDRs, including an insertion in one design and a deletion in the other. Furthermore, the two designs employ orthogonal edits, only sharing one position with different mutations (HC:S73R and HC:S73A) that is distant from the antigen and thus plausibly a neutral mutation. Folding of both sequences confirmed that the structural integrity of non-CDRH3 loops was maintained despite these changes (Supplementary Figure B14).

**Fig. 4:**
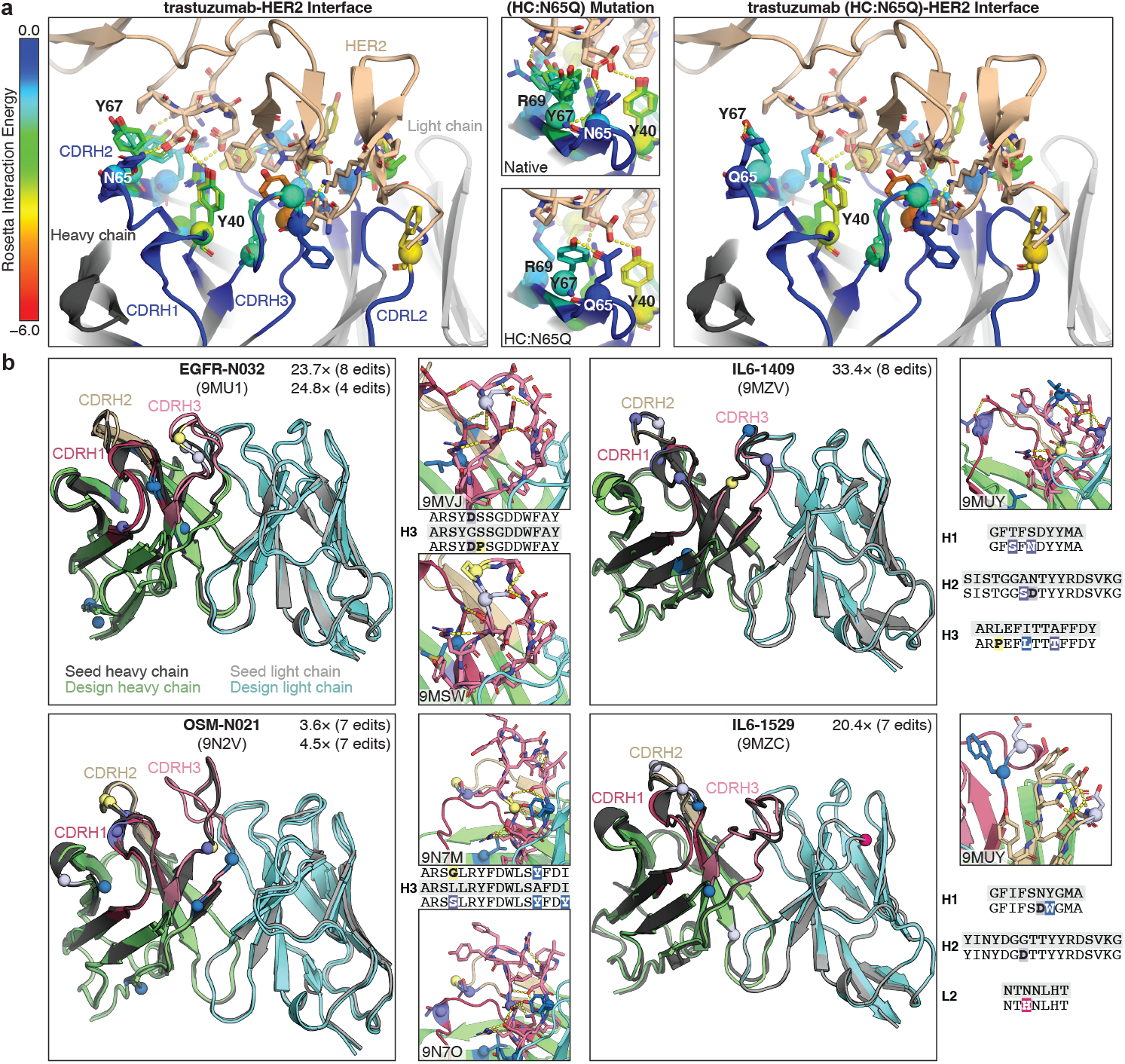
Interface structure modeling and experimental structure determination yield mechanistic insight into mutational effects. **a**, Native interactions from multiple replicate minimizations of trastuzumab (hu4D5) are shown as spheres and colored by Rosetta interaction energy (more negative is stronger), with hydrogen bonds depicted as yellow dashes. The CDRH2 interactions are highly dynamic, exhibiting multiple rotameric conformations with variable binding energy. Upon modeling the only mutation at a native contact from two separate *>*3× binders (HC:N65Q), a new hydrogen-bond between Q65-Y67 stabilizes the rotamers (highlighted in insets). All other mutations are separate from the native contacts (see Supplementary Figure B14), and also appear several places in the framework region, indicating that they act by indirect effects. **b**, Crystal structures of four seeds and six designs highlight very high structural consistency in almost all cases. For each design, the fold-increase in affinity and number of edits is specified. Edit positions are colored as: nonpolar (blue), polar (purple), negatively charged (lavender), positively charged (magenta), and conformational (yellow). The insets highlight regions of notable new interactions, primarily in CDRH3. Except for 9N7M, at least one edit in each design expands the intramolecular hydrogen-bond network within the CDRs. For the selected CDR sequences reported, the seed sequence is on a grey background and design sequences (insets) have mutations highlighted to match the renders.

This interface structure analysis suggests that our sequence-based design methods learn important structural and biophysical priors, resulting in optimized designs that preserve or optimize important energetic contacts necessary for binding, while introducing potentially epistatic mutations that improve binding affinity. Affinity maturation can occur via multiple routes [42] using distinct mutations that are not necessarily physically localized near the interface nor the antigen. We find that insertions and deletions are successfully introduced by our models and are tolerated, but not generally necessary for optimization.

### 2.6 Crystal structures of designed antibodies reveal mechanistic roles of mutations

To experimentally investigate the impact of designed mutations across a broader range of targets, we obtained apo crystal structures of four seeds and four designs (Supplementary Table B3, Supplementary Figure B15), in addition to two previously published (9MU1 and 9MSW [33]), spanning three unique targets (Fig 4**b**). SPR kinetic parameters and affinities are reported for all designs in this figure in Supplementary Table B4. For the non-OSM seeds, all crystallized designs are *>*10× improved binders. We observed remarkable conservation of overall CDR loop structure in 5/6 designs (consistent with predicted structures in Supplementary Figure B10), primarily through newly introduced hydrogen-bonding interactions (Fig 4**b** insets) that serve to further reinforce the seed loop conformations, which may contribute to affinity improvements via structural stabilization of binding-competent states even while unbound [43], though other mechanisms are possible [42]. Several other mutations make no new interactions with other residues (e.g. the mutations on CDRH2 of 9MUZ) but are highly solvent exposed and thus may constitute direct contacts to the antigen that are introduced without adversely affecting the native backbone.

The exception to this arose in the structural series for seed OSM-N021, where isosteric packing of G+Y mutations in the stem region of CDRH3 for design 9N7M (Δ*pK*_*D*_ = 0.56, 3.6× improvement) retained the conformations of all CDR loops, yet S+Y mutations at the same positions in design 9N7O (Δ*pK*_*D*_ = 0.65, nearly 4.5× improvement) produced an alternative loop conformation via novel hydrogen-bonding interactions that may contribute to its affinity maturation by more directly exposing the CDRH3 residues. The seeds OSM-N021, OSM-N088, IL6-951, and EGFR-N065 exhibit the least affinity improvement over all 4 rounds (Fig. 2b), but are also repeatedly measured among the highest absolute pKD values (Fig. 2d) and thus may already be near their local optima, consistent with our selection of viable lead candidates as starting points for optimization. It is therefore plausible that, in contrast to the native-contact-preserving edits to hu4D5 (Fig 4**a**) and the conformation-stabilizing edits to the other seeds (Fig 4**b**), the structural changes induced by edits in 9N7O may actually be critical to pushing affinity out of a local optimum imposed by the native backbone. The rarity of this occurrence is suggestive that structural consistency imposed by training data and filters (see “Biophysical Priors” in Section 4.1.2 of the Methods) is only sacrificed where more conservative mutations are insufficient.

## 3 Discussion

Here, we introduced the “Lab-in-the-loop” (LitL) system for online optimization of seed antibodies for multiple properties, including binding affinity and expression, against four distinct antigens of therapeutic interest: EGFR, OSM, HER2, and IL-6. LitL orchestrates generative and discriminative machine learning methods to explore sequence space, while remaining tightly coupled to high-quality, medium-throughput affinity and expression experimental feedback via active learning. Over four experimental iterations, we produced *>*3–100× higher binding affinity antibodies for ten seed molecules against all four different antigen targets, while simultaneously retaining or improving upon multiple other essential properties. Apo-state crystal structures of eleven seed antibodies and affinity matured designs (two previously published [33]) unambiguously demonstrated that LitL generates and selects designs possessing therapeutically relevant affinities (Table A1) while retaining the seed CDR loop conformations, in part through newly introduced rigidifying mutations and stabilizing intramolecular interactions. The models and filters described herein span both sequence and structural inputs, incorporating domain knowledge, biophysical priors, bioinformatics data, and model-driven *in vitro* data generation. We further demonstrate that sequence-based generative models can produce structurally interpretable mutations when a seed co-complex is available.

Lab-in-the-loop, as presented in this work, represents a framework by which end- to-end machine learning systems are developed and continually improved towards our ultimate goal of accelerating therapeutic antibody development via multi-property optimization. LitL is distinct from previous ML antibody maturation efforts (Section A) in that it is fundamentally designed around (and powered by) experimental antibody engineering priorities, constraints, capabilities, and challenges. Within this framework, experiments do not simply “close the loop” for validation of generated designs, but are instead embedded into all aspects of LitL. For example, the choice to ensemble models not only hedges against measurement noise and mitigates the limitations of any particular approach, it allows disparate families of models to be trained on highly diverse data modalities, producing unique generative and predictive capabilities. Similarly, by construction, selection can be facilely controlled both globally via biological priors and bioinformatic heuristics from decades of drug discovery experience as well as through bespoke specifications to address acute project needs. The advantages of this philosophy are evident in that, of the substantial improvements in antibody affinity reported herein, the best-performing molecules were obtained via orchestrating state-of-the-art ML modeling and antibody engineering approaches.

To comprehensively address the real-world challenges of therapeutic antibody development in practice, technical limitations to be addressed by future work include: improving sample and time efficiency to enable antibody optimization in fewer iterations and with greater accuracy, while preserving the modest scale of new requisite high-quality data acquired each cycle, and explicitly accounting for more numerous optimization objectives and constraints. Biological aspects of the problem, such as optimizing directly for biological function (not just affinity), and the early identification and triage of seed sequences by local optimality for more efficient resource allocation, are other active areas of research. Our modular lab-in-the-loop approach explicitly enables this evolution, demonstrating the power of active learning and algorithmic design of experiments.

Future work will unite these capabilities with parts of the drug discovery process beyond early stage discovery into an “end-to-end” machine learning system. By demonstrating acceleration of state-of-the-art experimental antibody engineering for therapeutic candidate antibodies and tackling the complexity of therapeutic discovery campaigns, we believe that LitL represents a significant step towards realizing the promise of ML and computation in large molecule drug discovery.

## 4 Methods

### 4.1 Computational methods

#### 4.1.1 Generative models

##### Unguided sampling for hit diversification

Unguided sampling for hit diversification is performed with discrete Walk-Jump Sampling (dWJS) and the Sequence-based Variational Diffusion Model (SeqVDM), introduced in Frey *et al*. [31]. dWJS is a generative model that combines separately trained score- and energy-based models (EBM) to learn a smoothed distribution of *noisy* data, sample noisy data from this smoother space with Langevin Markov Chain Monte Carlo (MCMC) and gradients from the EBM, and then denoise to “clean” data with the score-based model. dWJS builds on the Neural Empirical Bayes [44] formalism and generates 97-100% expressing and up to 70% functional, binding antibodies in *in vitro* experiments [31]. SeqVDM is adapted from [45] for protein sequences; it projects discrete sequences into a continuous latent space and performs denoising diffusion in the latent space. Further details for both dWJS and SeqVDM are provided in [31].

##### Guided sampling for lead optimization

Latent Multi-Objective Bayesian Optimization (LaMBO-2), introduced in Gruver & Stanton *et al*. [32], and Property Enhancer (PropEn) [34], are used for lead optimization via guided sampling. LaMBO-2 is a method for controlled categorical diffusion that enables sampling high likelihood, expressing protein sequences that are optimized with respect to an acquisition function, enabling multi-objective design. PropEn is a model designed to implicitly guide design optimization without the need for training a separate discriminator. Instead, PropEn leverages a matched training dataset. Initially, each sequence in the dataset is paired with a similar sequence that exhibits superior binding properties. A Seq2Seq model is then trained to take a lower affinity sequence as input and reconstruct its higher affinity counterpart. DyAb [33] leverages pairwise representations to predict property differences between antibody sequences. Here, it was employed as a ranking model to score combinations of known mutations, and as a generative model when combined with a genetic algorithm. More details regarding DyAb can be found in [33].

#### 4.1.2 Biophysical priors

Domain knowledge related to antibody engineering and protein design are incorporated into generative methods in the form of biological priors to restrict the high-dimensional search space and improve the rate of producing designs that bind to the antigen of interest. All the generative models used in this paper are *sequence-based* models; they are trained on amino acid sequences and emit amino acid sequences. While no protein structures are explicitly generated by our methods, protein structure information is included via strong biological priors. Models are trained on AHo-aligned [46] antibody sequences, a scheme with particular emphasis on structural consistency. AHo alignment introduces alignment tokens (“-”) and ensures a global sequence alignment across all antibodies, clearly separating an antibody sequence into distinct structural elements (framework and complementarity determining regions (CDRs)).

Training on aligned sequences [31, 32, 47, 48] is an established method for improving the performance of protein generative models and implicitly providing a strong structural prior. AHo alignment also enables our methods to easily introduce length changes (insertions or deletions), by sampling alignment tokens. However, at sampling time we often mask out the alignment token, preventing length changes, which may attenuate binding. Similarly, some designs restrict mutations to only the CDRs or subsets of the CDRs, which are generally thought to be most important for binding, though it is known that framework positions can also dramatically impact affinity, both individually [49, 50] and as part of humanization[51]. Our biophysical priors and design methods are built to preserve the binding mode of the seed antibody [52].

#### 4.1.3 Property prediction models

##### Multi-task fine-tuning

For each LitL round, denoted by *t*, we first perform *discriminative model selection*, using the generalization error on the last round (*t* − 1) to estimate the generalization error on the next round. A uniform random holdout set can be used as the validation set at *t* = 0. The best performing oracle is used for each property, as measured by a pre-determined metric on the hold-out set. Framing model selection as an open competition encourages rapid development of surrogate models and ensures that a diverse range of ML approaches are tried for challenging biological properties. An exceptionally performant model emerged during LitL competition: Cortex [32], a multi-task fine-tuning framework that uses pre-trained Language models for Biological Sequence Transformation and Evolutionary Representation (LBSTER) [53] to simultaneously model all properties of interest.

Lab-in-the-loop achieves better binding by 1) predicting antibody-antigen binding affinity, based on antigen and antibody sequence information, and selecting variants that are predicted to have improved affinity; and 2) by using generative models that either explicitly account for antigen information with the antigen sequence, or implicitly account for antigen information by using the antibody seed clonotype or training on matched pairs of sequences with improved binding.

##### Pre-trained representation learning

Language models for Biological Sequence Transformation and Evolutionary Representation (LBSTER) is used for pre-training representation learning models for multi-task property prediction and optimization with Cortex. LBSTER includes both Masked Language Models (MLMs) based on the ESM-2 model architecture [54] and Causal Language Models (CLMs) based on the Llama-2 model architecture [55]. LBSTER models are trained on Uniref50 [56] and the Observed Antibody Space [57], de-duplicated with MMSeqs2 [58]. We use the protein language model cramming strategy introduced in Frey *et al*. [53] to pre-train performative LBSTER models in only a few GPU days.

#### 4.1.4 Out-of-distribution detection

OOD detection is performed on designed sequences to determine if they are in-distribution for property prediction models and therefore can be reliably ranked by the ranking model. OOD detection is implemented with the distributional conformity score (DCS), as introduced in Frey *et al*. [31], which is a measure of how likely generated samples are with respect to a reference set of already known binders or expressors. DCS uses a likelihood under a joint density of statistical properties, including log-probability under a protein language model, and sequence-based properties like hydrophobicity and molecular weight, calculated with BioPython [59]. Kernel density estimation (KDE) is used to compute the joint density, although any density estimator may be used.

#### 4.1.5 Filters and quality control

Designs are annotated to ensure that they do not introduce chemical liabilities (e.g., glycosylation and deamidation motifs) and that conserved amino acids in antibody sequences are preserved (e.g., canonical cysteines). Additional quality control checks are performed to prevent germline reversion (which may lead to affinity attenuation). Further constraints are imposed across all designs and specific generative models to encourage generative models to explore local neighborhoods in sequence space around lead antibodies, so that designs are more likely to be in-distribution for property prediction models. In the first round of design, a maximum edit distance cap of 6 edits from the lead is enforced. In the second round, this cap is increased to 8, and in the third round, it is increased to 12. Some generative models restrict edits to particular regions (e.g., dWJS only introduces mutations in CDRs), while other methods including PropEn and LaMBO-2 introduce mutations across the entire sequence.

#### 4.1.6 Ranking and selection

At the beginning of each round, we draw a large pool of candidate designs from our generative models and pass them through the quality control filters. Because wet-lab assay measurements are time-consuming and expensive, only a small subset of the remaining candidates can be selected for experimental evaluation.

Formally, let *X* denote the design space of antibodies and let *f* : *χ* → R^*m*^ map a design **x** to a vector of *m* experimentally measured properties. The mapping *f* is treated as a *black box*: it can be queried but does not admit a closed-form expression. Our goal is to identify designs that approximate the *Pareto set*, i.e. the collection of *non-dominated* designs. A design **x**_1_ dominates **x**_2_ if it is no worse in every property and strictly better in at least one; a design is Pareto-optimal when no other candidate in *X* dominates it.

We tackle this problem with Bayesian optimization (BO), a sample-efficient framework for optimizing expensive black-box functions. Given labeled data 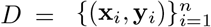, we fit a probabilistic surrogate model 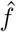 and rank new queries using an *acquisition function* 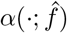 that balances exploitation (selecting designs predicted to be optimal) and exploration (selecting designs we are highly uncertain about). The top-scoring batch of designs under *ω* are synthesized, assayed, appended to *D*, and the cycle iterates.

Given a pessimistic reference point **r** ∈ ℝ^*m*^ (chosen to be strictly worse than the worst property values we are willing to accept), the *hypervolume* of a set *P* = {**y**_1_,…, **y**_|*P*|_} ⊂ ℝ^*m*^ is the hypervolume of the polytope dominated by *P* and bounded by **r** from below:

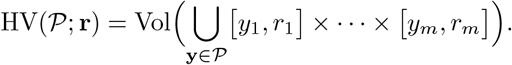

For a candidate design **x**^*′*^ with (unknown) property vector **y**^*′*^ = *f* (**x**^*′*^), its *hypervolume improvement* (HVI) with respect to the current batch *B* is

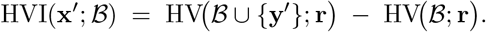

Because HVI measures the additional volume of property space dominated by **y**^*′*^, maximizing its expectation over the posterior *p*(*f* | *D*) (the expected HVI, or EHVI, acquisition function [60]) naturally favors designs that expand the Pareto front in different directions in the property space, thus promoting diversity among selected candidates.

In practice, the assay noise makes our observed property values random. We thus integrate over the uncertainty not only in the candidate designs but also in the observed designs, which yields the *noisy expected HVI* [NEHVI; 35]:

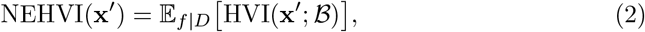

which provides robustness in noisy environments.

To form a batch of *k* designs in a given round, we employ the *sequential greedy* strategy: starting with *B*_0_ = ∅, we iteratively choose

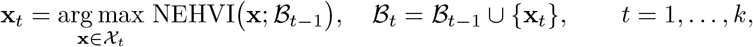

where *χ*_*t*_ is the pool of candidates in a given round *t*. This greedy procedure is computationally efficient, and empirical and theoretical results show that it is comparable to jointly optimizing *k* designs [61, 62].

Classical Bayesian optimization employs a Gaussian process (GP) posterior *p*(*f* | *D*) for surrogate modeling. Because LitL explores millions of high-dimensional sequence embeddings, GPs become computationally prohibitive. We therefore adopt a *deep ensemble* [63] of *S* independently initialized neural networks 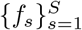. Each net-work outputs a predictive distribution over properties; treating each *f*_*s*_ as a Monte Carlo draw from the posterior yields an implicit approximation of *p*(*f* | *D*). We use these Monte Carlo draws to evaluate the expectation in Equation 2. Ensembling is a widely used strategy for capturing predictive uncertainty [64]: although any one model may underperform, the diverse, complementary strengths of its peers enable the ensemble to offset the shortcomings of individual networks.

#### 4.1.7 Interpretability

Sequence-level interpretability is performed with Cortex, as described in Adebayo *et al*. [65]. Independent Gaussian noise corruption is introduced as a computationally efficient alternative to adversarial training, without deteriorating classifier performance. Gaussian corruption is shown to confer faithfulness to feature attributions, i.e., saliency maps computed with classifiers that are regularized during training with Gaussian corruption correctly ignores spurious distractor features. This regularization allows inspection of gradient-based saliency maps to determine the residues and mutations that classifier predictions are most sensitive to.

#### 4.1.8 Rosetta biomolecular modeling

Input structures were prepared using Rosetta [39] gradient descent minimization followed by FastRelax [66, 67] with the *beta nov*16 scorefunction [40] and the following flags:

~~~
-corrections:beta_nov16 -ex1 -ex2 -use_input_sc -flip_HNQ -no_optH false -packing:repack_only true -relax:constrain_relax_to_start_coords -relax:ramp_constraints false
~~~

In the case of the seed hu4D5 (trastuzumab), coordinates were obtained from PDB code 1n8z. The HC:N65Q mutation was introduced using the MutateResidue mover. The entire structure was first minimized using MinMover then the interface was allowed to repack using FastRelax (InterfaceRelax2019 script) with nonramping backbone constraints to the starting coordinates applied during both steps. This MinMover+FastRelax protocol was found to produce plausible local variation in sampling more effectively than either Mover alone, while still yielding low-energy structures expected of a high-affinity protein complex. Two-body interaction energies were computed using the InteractionEnergyMetric, which includes through-space terms including dispersion, solvation, electrostatic, and hydrogen bonding interactions. Contacts were defined as any residue that has ≤ −0.6 REU interaction energy to at least one individual residue on the antigen side. Per-residue values were written to the B-factor columns and used for visualization. All other visualized structures were folded directly from the seed or design sequences using ABodyBuilder2 and saved with refinement.

### 4.2 Engineering infrastructure

#### 4.2.1 Data engineering

Our data backend serves as an intermediary connecting functional assay data curated by scientists, direct instrument access, as well as data drawn from public and proprietary databases. In the context of LitL, this constitutes the retrieval and processing of assay data to enable retraining and validation of our models. At a high level, we implemented abstractions for periodic (i) access, (ii) retrieval, and (iii) transformation of data to produce and update core datasets. Together this enables database agnostic deployable pipeline orchestration that persists dataset artifacts in the cloud, providing a common setting in which data from disparate sources may be jointly queried, combined, quality controlled, and ultimately packaged into ML-ready datasets.

In addition to basic data integrity, reproducibility, and versioning carried out automatically by data pipelines, quality control standardizes assay registration and data capture. We set protocols for instrument runs, machine data capture, as well as the application of human and heuristic annotation. As a result of these efforts, we have dramatically increased available data across many more features and with sufficient assay context to drive model improvement.

Designed sequence libraries are verified and registered in the internal large molecule registration system, and each sequence receives a unique identifier. Protein production requests at a scale of 1 ml are submitted via an internal protein production system, and all related request and fulfillment metadata are collected into databases and linked with unique identifiers from the registration system. Subsequent functional assays are recorded in a structured fashion via an internal Lab Information Management System (LIMS) that also supports plating provenance and registrations across all manual and automated Antibody Engineering workflows. Data for each molecule is centralized and integrated into a single repository (Supplementary Figure B16) and made available for consumption by downstream machine learning workflows.

### 4.3 Design submission

When a design is submitted, metrics to quantify binding affinity and expression yield predictions are computed. Litl is orchestrated using a Kubernetes-based computation pipeline, providing a structured and modular approach to pipeline development. The frontend user interface manages instructions for file submission, file upload to persistent storage, and visualizations of the uploaded antibody data. Alternatively, a Python client can be used for programmatic submission and retrieval of pipeline results. Following file upload, the server performs data formatting checks and triggers the computation pipeline, which executes antibody registration, quality checks, enumeration, folding, inference with property prediction models, out-of-distribution checks, and quality control filtering. Parallel execution is employed for independent tasks. Processed data undergo additional quality control before being passed to the ranking pipeline, using an active learning framework. The selection of the best candidate designs is then passed into an automated visualization tool and a formatting step for DNA synthesis order.

### 4.4 Experimental methods

Experimental protocols follow Hsiao *et al*. [4, 68] and Wu *et al*. [36]. The methods are described here for convenience.

#### 4.4.1 Immunization

We follow the animal immunization procedure described in Hsiao et al. [4]. Animals were immunized and both splenocytes and lymph node cells were used for hybridoma fusion. RNeasy Mini Kit (Qiagen) was used for total RNA extraction from hybridoma cells. Reverse-transcription (RT) PCR of heavy and light chains was done with a SMARTer RACE cDNA Amplification Kit (Clontech). The forward Universal Primer Mix from SMARTer RACE cDNA Amplification Kit was used for the PCR step. The dideoxy sequencing method was used to directly sequence amplified PCR products.

#### 4.4.2 Repertoire sequencing and mining

Direct lysis with ammonium chloride was used to remove red blood cells. Spleen and bone marrow cells were also collected. Buffer RLT (Qiagen) was used to lyse remaining splenocytes and bone marrow cells. The RNeasy Mini Kit was used to purify RNA according to the instructions from the manufacturer. Amplicons were purified from agarose gels and sequenced with an Illumina MiSeq instrument. Sequences were processed to 1) extract sequences within variable region boundaries, 2) remove incomplete sequences and sequences with ambiguous residues, 3) remove the first eight residues of framework region 1 (FR1) encoded by amplification primers, 4) extract sequences matching mAb CDR3 length and identity threshold of 57% for CDR H3 sequences, and 5) identify and count unique variable region sequences [4].

To compare against design libraries from generative models, some heavy chain variants of seeds (with light chain fixed) were either chosen directly from the bulk NGS data or designed based on observed patterns in the neighborhood of the seed sequences. First, we collected all sequences that have the same lengths as seeds and are within 12 Hamming distance over the entire variable region and 3 Hamming distance in CDR-H3 from the seeds. Among these, we picked several observed sequences that are most distant by Hamming distance from inferred germline sequences associated with seed sequences. Next, we used the collected sequences to construct Position Specific Scoring Matrix (PSSM) including each seed and their neighbors. We then designed one sequence per seed by sampling the frequency mode of the PSSM at each position while preventing reversion to inferred germline sequence residues associated with the seed. We also included observed sequences that rank high according to the PSSM (i.e. sum of frequencies of observed residues).

#### 4.4.3 Plasmid construction and antibody production

All antibodies were expressed utilizing GLFs as previously described [36]. Briefly, VH and VL fragments were synthesized with 40bp overhang sequences on both sides (Twist Bioscience) and subsequently integrated via Gibson assembly into in-house designed pRK5 expression vectors containing the appropriate heavy chain or light chain constant regions and other elements required for expression (promoter, ORF, poly-A tail) [69]. The product of this reaction was used as the template for GLF amplification, with primers located back-to-back in the noncoding region of the vector, amplifying the whole vector-sized linear DNA insert. This insert contains 2kb extra sequence on each side of the expression cassette, offering extra protection from degradation.

1mL scale IgG expressions were transiently conducted in Expi293F cells (ThermoFisher Scientific). Cells were transfected using PEIpro (PolyPlus PEIpro, Reference #: 115-01L. 1mg/ml). 1µL PEIpro was diluted into 100µL serum-free medium, then 6µL of GLFs (1:1 mixture of heavy chain and light chain, approximate concentration 200µg/µL) were added and incubated for 10 min. The PEI-DNA complex was added to the 0.85ml cells, and cells were cultured in a 96-deep well plate at 37 °C and 8% CO2 for 7 days, with 150µL fresh medium added 24hr post transfection.

The supernatant was collected and purified with a 1-step purification using Protein A affinity resin (MabSelect SuRe™, Cytiva). Quality control of antibody purity was determined by SDS-PAGE with Coomassie blue staining and A280 absorbance to measure protein concentration.

#### 4.4.4 SPR affinity characterization

Surface Plasmon Resonance (SPR) was performed on a Biacore 8K+ (Cytiva). The instrument was primed with HBS-EP buffer (0.01 M HEPES pH 7.4, 0.15 M NaCl, 3 mM EDTA, 0.005% v/v Surfactant P20), and antigens or antibodies were also diluted into the same buffer for all SPR experiments. A Protein A chip was used for antibody capture in all experiments. Experiments were either done using single cycle kinetics (the antigen series successively injected from low to high concentrations, followed by one dissociation step) or multi cycle kinetics (each cycle using one antigen concentration for association followed by a dissociation step, and then repeated for each antigen concentration). In a typical experiment, five concentrations (0.16nM, 0.8nM, 4nM, 20nM and 100nM) for 180s association time, 600s dissociation time with a Flow rate of 30 µl/min. All data were analyzed using Biacore Insight software using a 1:1 binding model.

#### 4.4.5 BV ELISA

BV ELISA was performed following [38], using chimeric mouse/human IgG1 antibodies. Antibodies were purified by protein A affinity chromatography and SEC. BV ELISA is an assay based on ELISA detection of non-specific binders to baculovirus (BV) particles. BV ELISA identifies antibodies that have an increased risk for poor pharmacokinetic properties (fast clearance). Molecules with a BV ELISA score *>* 1.0 indicate an increased risk. We train a surrogate model by fine-tuning a pre-trained LBSTER [53] protein language model on an internally generated dataset of 2,305 BV ELISA measurements.

#### 4.4.6 Protein expression, purification, and structure determination

The Fab constructs were transfected into CHO cells at a 1:2 (HC:LC) DNA ratio, and expressed for 10 days. After harvesting, the supernatant was collected for purification using GammaBind Plus Sepharose (Cytiva) followed by size exclusion or SP cation exchange chromatography. The purified Fabs were formulated in 20mM histidine acetate pH 5.5, 150mM sodium chloride.

The Fabs were concentrated to 10mg/mL and crystallized by vapor diffusion as follows: 9MVJ: 2.0 M Ammonium sulfate, 0.2 M Sodium chloride, 0.1 M Sodium cacodylate pH 6.5; 9MUZ: 0.2 M Sodium Potassium phosphate, 20% v/v Polyethylene glycol 3350; 9MUY: 0.1 M Ammonium acetate, 0.1 M Zinc chloride, 0.1 M Bis-Tris pH 7.2, 15% v/v PEG Smear High; 9MZC: 0.09 M NPS, 0.1 M BS1 pH6.5, 50% v/v, PM2 (Polymer); 9MZV: 0.2 M Calcium chloride, 0.1 M Tris-HCl pH 8.5, 20% w/v PEG 4000; 9N2V: 0.2 M Sodium chloride, 0.1 M Imidazole pH 8.0, 0.4 M NaH_2_PO_4_, 1.6 M K_2_HPO_4_; 9N7M: 0.2 M Ammonium sulfate, 0.1M MES pH 6.5, 30% w/v PEG 5000 MME; 9N7O: 0.1 M Bis-Tris propane pH 9.0, 10% v/v PEG 200, 18% w/v PEG 8000;

Crystals were cryoprotected by addition of ethylene glycol. Data were collected at synchrotron sources, as indicated in Supplementary Table B3, and data were scaled using XDS [70]. The structures were determined by molecular replacement, using PHASER [71], and the models were built in COOT [72] and subsequently refined in PHENIX [73] to final statistics presented in Supplementary Table B3. The simulated annealing omit difference electron density maps (Supplementary Figure B15) demonstrate the fidelity of the final models with the electron density.

## Supporting information

Appendix and References

## Supplementary information

Supplementary files accompany this article.

## Acknowledgments

We are grateful to Viva Biotech staff, Yanshu Dou, Chunheng Huo, Yu Tang for contributions to the crystal structures and to Feng Wang and Yoana Dimitrova, for coordinating structural experiments. The authors would like to thank the facilities and staff of Diamond Light Source beamline I03 and Shanghai Synchrotron Radiation Facility (SSRF) beamline BL02U1 for contributions towards crystallographic data collection. We thank the experimental facility and the technical services provided by “The Synchrotron Radiation Protein Crystallography Core Facility (SPXF) of the National Core Facility for Biopharmaceuticals (NCFB), National Science and Technology Council (NSTC)” and the “National Synchrotron Radiation Research Center (NSRRC)”, a national user facility supported by the National Science and Technology Council of Taiwan (NSTC), Taiwan (R.O.C.). We acknowledge DESY (Hamburg, Germany), a member of the Helmholtz Association HGF, for the provision of experimental facilities. Parts of this research were carried out at beamline P11.

We thank Jia Wu, Kellen Schneider, and Dhaya Seshasayee for the in vivo antibody discovery work. We also thank the Genentech NGS core lab for the repertoire sequence data used in this study.

The authors acknowledge the entire Prescient Design team and the Antibody Engineering department at Genentech for providing helpful discussions and input that contributed to the research results reported within this paper. The authors would especially like to acknowledge Candice Wertz for coordination and project management for manuscript preparation.

## Declarations

### Ethics approval

All animal studies were performed in animal facilities accredited by the Association for Assessment and Accreditation of Laboratory Animal Care International. The procedures for animal studies were compliant under the Institutional Animal Care and Use Committee of the facility.

### Availability of data and materials

LBSTER models were trained using the UniRef50, PDB, and Observed Antibody Space databases. All other models were trained on internally generated datasets described in the experimental methods, Section 4.4.

Crystallographic coordinates and data were deposited at the RCSB with accession codes 9MUY, 9MUZ, 9MVJ, 9MZC, 9MZV, 9N2V, 9N7M, 9N7O.

### Code availability

Discrete Walk-Jump Sampling, PropEn, LBSTER, and Cortex are available on the Prescient Design GitHub. DyAb is available as a model within LBSTER. The Prescient standard library (Beignet) and LaMBO-2 are available on GitHub.

### Authors’ contributions

R.G.A., D.B., R.B., K.C., H.D., N.C.F., V.G., I.H., R.K., S.K., J.R.K., J.T.K., J.LV., J.H.L., A.L., S.R., F.S., S.D.S., and A.M.W. conceived of and designed the work. R.G.A., J.B., T.B., P.C., Y.C., T.D., H.D., A.D., J.F., N.C.F., V.G., A.G., J.H., H.I., A.I., S.J., S.K., M.K., J.K., J.L.V., A.L.F., E.L., W.C.L., J.Y.Y.L., S.P.M., E.K.M., S.N., J.W.P., J.P., S.R., F.S., S.D.S., N.T., H.T., A.W., A.M.W., B.W., S.W., and K.Z. acquired the data. R.G.A., D.B., J.B., T.B., P.C., Y.C., A.C., T.D., H.D., N.C.F., V.G., A.G., I.H., H.I., T.J., S.K., J.R.K., J.K., J.L.F., J.H.L., E.L., D.L., W.C.L., A.L., J.L., O.M., E.K.M., K.M., H.M.P., J.P., S.R., S.R., F.S., I.S., A.S., S.D.S., E.W., A.W., A.M.W., B.W., and K.Z. analyzed and interpreted the data. R.G.A., D.B., T.B., H.D., N.C.F., V.G., A.G., J.H., H.I., A.I., S.K., M.K., J.K., A.L.F., E.L., J.Y.Y.L., A.L., J.L., S.P.M., O.M., S.N., J.W.P., J.P., S.R., S.S., S.D.S., N.T., E.W., A.M.W., and K.Z. created methods and algorithms presented in this work. R.G.A., R.B., K.C., A.D., J.F., N.C.F., V.G., J.H., I.H., S.J., T.J., R.K., S.K., J.R.K., J.T.K., J.L.F., J.H.L., A.L., E.K.M., J.M., K.M., H.M.P., J.W.P., S.R., A.R., S.R., S.S., I.S., S.D.S., N.T., A.W., A.M.W., and Y.W. wrote and edited the manuscript. R.B., P.C., Y.C., K.C., H.D., V.G., I.H., R.K., E.K.M., S.K., J.R.K., J.T.K., J.L.V., A.L., J.M., S.R., A.R., F.S., A.M.W., B.W., and Y.W. supervised the research. All authors discussed the results and approved the final version of the manuscript.

