## Appendix and References for "Lab-in-the-loop therapeutic antibody design with deep learning"

#### A Supplementary evaluations and comparisons to prior work

We conduct comparisons relating our work and results to previously published anti-(OSM, EGFR, HER2, IL6) antibodies as well as prior protein design work in the literature. Although a head-to-head comparison is not possible, because our antibodies and other published antibodies differ in targeted epitope, function, and germline, we contextualize our results in terms of other previously published antibodies. The antibodies reported in our work are comparable to or surpass binding affinities of those reported previously, while also optimizing for expression yield and developability constraints.

Lab-in-the-loop is a system for designing experiments and executing multi-property optimization of antibodies in the drug discovery context, rather than a single model or component like EffEvo [16] or EVOLVEpro [15], and is not related to *de novo* design methods like RFDiffusion. Significant benchmarking of LitL's constituent models has already been reported within their primary publications (see associated references within the Methods), which we summarize below for convenience.

We have extensively benchmarked the performance of various protein language models (AntiBERTy, ESM2, and our own LBSTER language models) [33] for predicting binding affinities. We found that no language model embedding strictly dominated the others across antigen targets and antibody leads/seeds, which contributed to our present strategy of ensembling over methods. See Ref. [33] for further details.

In *in silico* benchmarks optimizing sequences with both guided and unguided generative methods [31, 32], we found that our methods outperformed RFDiffusion, DiffAb, DiGress, IgLM, ESM2, and genetic algorithms (AdaLead, PEX) in terms of optimizing objective values while preserving antibody sequence likelihood. In a comparison to AbSci's [74] originally published results, our models produced higher rates of binding and rates of improved binding, while also respecting naturalness (likelihood) constraints. See Refs. [31, 32] for further details.

##### A.1 ML protein design baselines from prior work

Although it is not typical among the referenced prior works to compare against other ML methods, for completeness we now report 360 antibody variants targeting HER2, OSM, IL6, and EGFR generated using two previously published approaches: EffEvo [16] and ProteinMPNN [75]. To make these baselines more competitive and increase the informational value of the resulting experimental data, we tuned and adapted them for Lab-in-the-Loop. We tuned ProteinMPNN sampling parameters to optimize for antibody CDR sequence recovery rate, implemented EffEvo (using the original suite of language models as well as AntiBERTy [19]), and additionally devised a structurally informed variation (ESM-Spatial-Rosetta) that uses predicted antibody

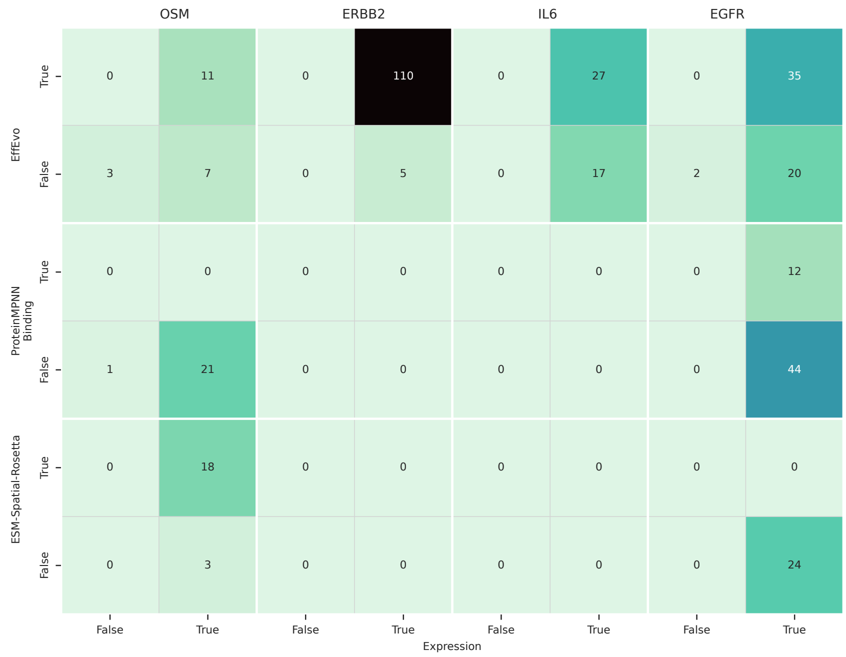


**Fig. A1**: Expression and binding counts for all targets across three generative ML baselines.

structures and PyRosetta to identify favorable combinations of mutations. These baselines were performed prior to Round 1 across a large number of seeds, and thus capture general trends representing the ability of these models to perform complex therapeutic optimization from many starting sequences in the absence of LitL-informed data points.

Under global selection, all baseline methods generated >97% expressing antibody variants and individually exhibited biased suitability for different targets (Fig. A1), a property we similarly noted above that motivates the choice of model ensembling to improve target generality. EffEvo has a limited edit distance window (mean edit distance of 3.86), which preserves binding at a high rate of 77%, but is sample inefficient for exploring the mutational space (Fig. A2). We attempted to strengthen the method's performance using PyRosetta and antibody structure prediction (ESM-Spatial-Rosetta), which preserved binding in 40% of designs, but the fundamental limitation of small edit distance windows, and thus sample inefficiency, remained. We have previously attempted to increase the edit distance neighborhood of EffEvo [31], but because the underlying method is a pre-trained ESM2 BERT-style protein language model trained at a constant 15% masking rate, it struggles to preserve or improve binding at higher edit distances relevant for efficient optimization loops.

We additionally benchmarked ProteinMPNN, including tuning the sampling temperature to maximize sequence recovery rate on internal datasets of antibody sequences. We predicted seed antibody structures using ABodyBuilder2 [76] and

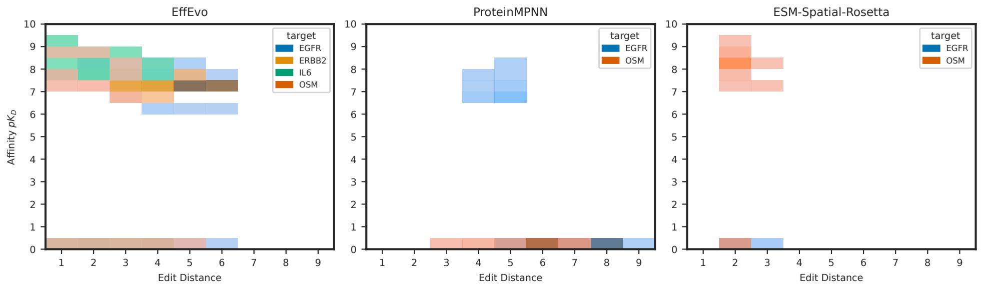


**Fig. A2**: 2D histogram of affinity $pK_d$ versus mutational load for designs generated by external sequence- and structure-based ML baselines.

used ProteinMPNN for sequence diversification. ProteinMPNN effectively designs 3-9 residues (mean 6.33) simultaneously in the CDRs, a larger number than EffEvo or Spatial Rosetta EffEvo, but the binding rate of ProteinMPNN designs was 15%, much lower than the other generative models used in our work and only successful for the EGFR target (Fig. A2).

Most importantly, all baseline methods tend to shift the pKD distribution to lower affinities, rather than higher, with increasing numbers of mutations. No binding was observed for any design with >6 mutations across all targets, in contrast with edit distances of up to 14 from LitL's suite of generative models (Figs. B11 and B3). We hypothesize that this behavior arises because the methods are trained on general protein families (Uniref or PDB) and tend to revert antibody sequences closer to germline, which attenuates affinity.

#### A.2 Comparison to previously published antibodies

Antibodies presented in this work possess affinities comparable to literature values reported for clinical antibodies against EGFR, OSM, and HER2, and weaker binders than those previously reported for IL6. These antibodies differ in targeted epitope, function, germline, and binding mode, but a comparison is given here for completeness. Table A1 below summarizes the results.

Published anti-EGFR antibodies include cetuximab (200 pM) [77], panitumumab (50 pM) [78], and GC1118 (160 pM) [79], compared to the best anti-EGFR design presented in this work at 47 pM binding affinity.

We have previously reported antibody designs with similar or better binding affinities to anti-HER2 antibodies, including trastuzumab and pertuzumab [31]. Relative to another ML-guided optimization of trastuzumab [80], we obtain a further 3-fold improvement in relative affinity increase compared to their best clone.

There are few reported anti-OSM antibodies; the best reported in this work (291 pM) is of similar affinity to GSK2330811, measured to be 568 nM *in vivo* [81].

Finally, though the best reported anti-IL6 antibodies in this work are weaker binders than olokizumab (10 pM) [82] and 61H7 (3 pM) [83], the highest affinity measured (197 pM) is still well within typical sub-nM therapeutic values.

**Table A1**: LitL designs compared to public therapeutic candidate antibodies.

| Target | Highest affinity design | Fold-change from seed | Comparator molecule | Comparator affinity |
|---|---|---|---|---|
| EGFR | 47 pM | 39.8× | panitumumab | 50 pM |
| ERBB2[a] | 206 pM | 9.1× | Mason et al. 2021 | 140 pM (2.9×) |
| IL6 | 197 pM | 106.6× | olokizumab | 10 pM |
| OSM | 291 pM | 6.7× | GSK2330811 | 568 pM |

[a] Absolute measured affinity of hu4D5 by antibody-capture SPR as described in Methods 4.4 can be highly variable. Fold-change normalizes against these batch effects. The seed affinity reported in Mason et al. of 400 pM is close to the true value.

903
### B Supplementary data

**Table B2**: SPR binding kinetics for select antibody designs reported in the main paper. Full antibody and antigen sequences and additional data, including goodness of fit metrics, are available in the Extended Data.

| Antibody Alias | $k_a$ (1/Ms) | $k_d$ (1/s) | $K_D$ (M) | pKD | Target |
|---|---|---|---|---|---|
| EGFR-P01451 | 5.29e+06 | 4.50e-04 | 8.50e-11 | 1.01e+01 | EGFR |
| EGFR-P01484 | 4.12e+06 | 6.16e-04 | 1.50e-10 | 9.82e+00 | EGFR |
| EGFR-P01503 | 3.99e+06 | 6.06e-04 | 1.52e-10 | 9.82e+00 | EGFR |
| EGFR-P01519 | 2.16e+06 | 1.44e-02 | 6.68e-09 | 8.18e+00 | EGFR |
| EGFR-P01606 | 3.64e+06 | 5.96e-04 | 1.64e-10 | 9.79e+00 | EGFR |
| ERBB2-P00873 | 1.20e+05 | 2.01e-04 | 1.67e-09 | 8.78e+00 | ERBB2 |
| ERBB2-P00961 | 1.25e+05 | 3.14e-04 | 2.52e-09 | 8.60e+00 | ERBB2 |
| ERBB2-P00965 | 1.22e+05 | 2.68e-04 | 2.20e-09 | 8.66e+00 | ERBB2 |
| IL6-P01545 | 1.27e+06 | 1.17e-03 | 9.18e-10 | 9.04e+00 | IL6 |
| IL6-P01548 | 8.70e+05 | 8.18e-04 | 9.41e-10 | 9.03e+00 | IL6 |
| IL6-P01573 | 2.20e+06 | 1.19e-03 | 5.40e-10 | 9.27e+00 | IL6 |
| IL6-P01577 | 1.13e+06 | 1.50e-03 | 1.33e-09 | 8.88e+00 | IL6 |
| IL6-P01615 | 6.59e+05 | 9.64e-04 | 1.46e-09 | 8.84e+00 | IL6 |
| IL6-P01634 | 5.82e+05 | 3.53e-04 | 6.07e-10 | 9.22e+00 | IL6 |
| IL6-P01662 | 9.76e+05 | 1.37e-03 | 1.41e-09 | 8.85e+00 | IL6 |
| IL6-P01770 | 1.16e+06 | 1.11e-03 | 9.61e-10 | 9.02e+00 | IL6 |
| OSM-P01123 | 1.13e+06 | 1.83e-03 | 1.61e-09 | 8.79e+00 | OSM |
| OSM-P01250 | 6.64e+05 | 3.55e-04 | 5.34e-10 | 9.27e+00 | OSM |
| OSM-P01344 | 3.67e+05 | 3.55e-04 | 9.67e-10 | 9.01e+00 | OSM |
| OSM-P01377 | 1.04e+06 | 1.72e-03 | 1.66e-09 | 8.78e+00 | OSM |
| OSM-P01473 | 3.31e+05 | 2.69e-04 | 8.12e-10 | 9.09e+00 | OSM |
| OSM-P01508 | 1.30e+05 | 3.32e-04 | 2.55e-09 | 8.59e+00 | OSM |

**Table B3**: Crystallographic data

| PDB code | 9MVJ | 9MUZ | 9MUY | 9MZC |
|---|---|---|---|---|
| X-ray source | NSRRC TPS 05A | SSRF BL02U1 | Diamond I03 | SSRF BL02U1 |
| Wavelength (Å) | 0.99987 | 0.97918 | 0.97625 | 0.97918 |
| Detector | Rayonix MX300HS | Eiger2 S 9M | Eiger2 XE 16M | Eiger2 S 9M |
| Resolution range (Å) | 49.3 – 1.95 | 147.3 – 1.71 | 50.9 – 1.97 | 47.5 – 1.58 |
| Highest res. bin (Å) | 2.00 – 1.95 | 1.74 – 1.71 | 2.02 –1.97 | 1.60 – 1.58 |
| Space group | $H3_2$ | $P2_12_12$ | $C_2$ | $P2_12_12$ |
| Multiplicity | 20.7, 21.2 | 7.0, 6.0 | 6.9, 6.6 | 6.9, 7.1 |
| Complete (%) | 100, 100 | 100, 100 | 99.9, 99.8 | 98.1, 95.1 |
| Mean $I/\sigma_I$ | 21.0, 2.5 | 11.3, 1.9 | 13.3, 2.2 | 12.4, 2.0 |
| Wilson B (Å$^2$) | 33.8 | 19.6 | 32.1 | 3.5 |
| CC ½ (highest bin) | 0.86 | 0.69 | 0.87 | 0.69 |
| Rmerge (%) | 10.7, 159 | 10.8, 86.1 | 7.7, 80.9 | 16.7, 107 |
| # reflections (Rfree set) | 59,055 (5,735) | 91,776 (8,797) | 38,938 (3,770) | 62,385 (3,054) |
| Resolution range (Å) | 39.7 – 1.95 | 73.7 – 1.71 | 33.3 – 1.97 | 47.3 – 1.58 |
| Rwork, Rfree (%) | 18.0, 19.8 | 16.6, 19.6 | 20.8, 22.3 | 17.5, 20.2 |
| # non-H atoms | 3,715 | 7,682 | 3,585 | 4,264 |
| # solvent molecules | 328 | 939 | 225 | 606 |
| Rmsd bond lengths (Å) | 0.004 | 0.004 | 0.005 | 0.006 |
| Rmsd bond angles (°) | 0.668 | 0.644 | 0.832 | 0.913 |
| Ramachandran fav. (%) | 97.4 | 98.1 | 96.5 | 97.9 |
| Ramachand. outlier (%) | 0 | 0 | 0.24 | 0 |
| Ave B-factor (Å$^2$) | 48.2 | 23.3 | 53.9 | 12.6 |
| Molprobity clash score | 1.82 | 1.43 | 3.97 | 1.67 |

| PDB code | 9MZV | 9N2V | 9N7M | 9N7O |
|---|---|---|---|---|
| X-ray source | PETRAIII P11 | PETRAIII P11 | SSRF BL02U1 | SSRF BL02U1 |
| Wavelength (Å) | 1.03321 | 1.03321 | 0.97918 | 0.97918 |
| Detector | EIGER2 X 16M | EIGER2 X 16M | Eiger2 S 9M | Eiger2 S 9M |
| Resolution range (Å) | 48.3 – 1.90 | 46.55 – 1.97 | 46.53 – 2.15 | 44.9 – 1.48 |
| Highest res. bin (Å) | 1.94 – 1.90 | 2.00 – 1.97 | 2.20 – 2.15 | 1.50 – 1.48 |
| Space group | $P2_1$ | $P4_1$ | $P4_1$ | $P4_12_12$ |
| Multiplicity | 6.9, 6.9 | 13.7, 13.4 | 5.1, 4.2 | 26.1, 21.1 |
| Complete (%) | 99.3, 99.1 | 99.5, 100 | 99.9, 100 | 99.8, 100 |
| Mean $I/\sigma_I$ | 12.4, 2.2 | 16.1, 2.1 | 7.7, 2.0 | 13.0, 2.0 |
| Wilson B (Å$^2$) | 20.6 | 27.8 | 19.0 | 19.6 |
| CC ½ (highest bin) | 0.68 | 0.745 | 0.154 | 0.306 |
| Rmerge (%) | 13.4, 103 | 15.4, 162 | 22.4, 70.7 | 18.2, 184 |
| # reflections (Rfree set) | 71,330 (3,484) | 95,351 (4,880) | 73,985 (3,682) | 94,027 (4,649) |
| Resolution range (Å) | 48.3 – 1.90 | 46.6 – 1.97 | 41.3 – 2.15 | 31.8 – 1.48 |
| Rwork, Rfree (%) | 17.7, 21.4 | 17.4, 20.2 | 21.0, 24.2 | 17.2, 20.5 |
| # non-H atoms | 7,507 | 7,635 | 7,474 | 3,500 |
| # solvent molecules | 839 | 853 | 826 | 601 |
| Rmsd bond lengths (Å) | 0.002 | 0.002 | 0.002 | 0.006 |
| Rmsd bond angles (°) | 0.591 | 0.589 | 0.512 | 0.841 |
| Ramachandran fav. (%) | 98.0 | 98.0 | 98.1 | 97.7 |
| Ramachand. outlier (%) | 0 | 0 | 0 | 0 |
| Ave B-factor (Å$^2$) | 24.0 | 38.0 | 27.5 | 26.1 |
| Molprobity clash score | 0.92 | 0.82 | 1.15 | 1.29 |

**Table B4**: SPR binding kinetics for designs reported in Figure 4. Full antibody and antigen sequences and additional data, including goodness of fit metrics, are available in the Extended Data. ΔpKD values are reported for measurements where design and seed were on the same plate, leading to two slightly different affinities for the OSM-N021 seed. "N/A" PDBIDs are the two folded anti-HER2 designs for which experimental structures were not obtained. Affinities for 9MSW and 9MU1 were previously reported in [33].

| PDBID | Antibody Alias | Round | $k_a$ (1/Ms) | $k_d$ (1/s) | $K_D$ (M) | pKD | Seed pKD | Δ pKD |
|---|---|---|---|---|---|---|---|---|
| 9MVJ | EGFR-P01451 | 3 | 4.86E+06 | 6.24E-04 | 1.28E-10 | 9.891 | 8.517 | 1.374 |
| 9MSW | EGFR-P01550 | 3 | 4.60E+06 | 5.65E-04 | 1.23E-10 | 9.911 | 8.517 | 1.394 |
| 9N7M | OSM-P01295 | 2 | 2.78E+05 | 1.30E-04 | 4.68E-10 | 9.330 | 8.770 | 0.560 |
| 9N7O | OSM-P01513 | 3 | 3.26E+05 | 1.41E-04 | 4.33E-10 | 9.364 | 8.708 | 0.656 |
| 9MUZ | IL6-P01770 | 3 | 1.67E+06 | 1.03E-03 | 6.17E-10 | 9.210 | 7.686 | 1.524 |
| 9MUY | IL6-P01634 | 3 | 2.24E+06 | 3.28E-04 | 1.46E-10 | 9.834 | 8.487 | 1.347 |
| N/A | ERBB2-P00873 | 3 | 1.28E+05 | 1.67E-04 | 1.30E-09 | 8.884 | 8.195 | 0.689 |
| N/A | ERBB2-P00965 | 3 | 1.39E+05 | 2.04E-04 | 1.47E-09 | 8.833 | 8.195 | 0.638 |

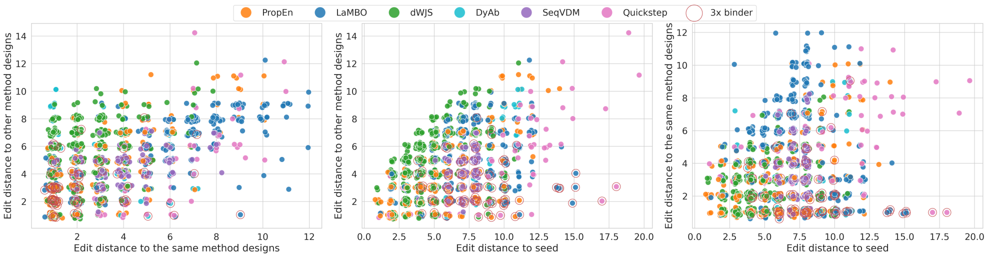


**Fig. B3**: Edit distances of generated designs to seed, other method designs, and other designs from the same method. It can be seen that 3x binders usually have lower edit distances to other designs within and between generative methods, but *higher* edit distances to seed. This highlights an important aspect of our multi-round approach, namely, the ability to move further from seed in a productive manner. It is also worth noting that while most methods have rather linear relationship between distance to seed and distance to other method designs, with a slightly higher similarity to other method designs, dWJS can produce tangent designs as shown by low edit distance to seed and high edit distance to designs from other methods.

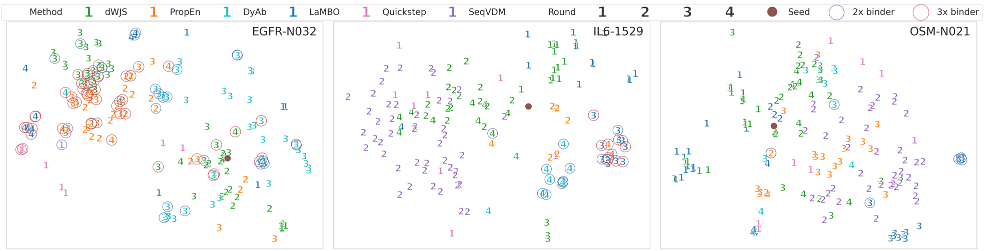


**Fig. B4**: t-SNE[84] plot of ESM-2 [54] embeddings of designs around 3 selected seeds. We can see that dWJS and SeqVDM, as expected, explore the design space more widely, while designs of PropEn and LaMBO are more localized, especially within the same round. The importance of thoroughly exploring the space around the seed is also highlighted by the fact, that most of the time 2x and 3x binders are tightly grouped and can be far from the seed. The Quickstep designs hand-picked from NGS sometimes, for some seeds, can lead to 3x binders, but for many seeds they do not explore in the regions holding the 2x and 3x binders.

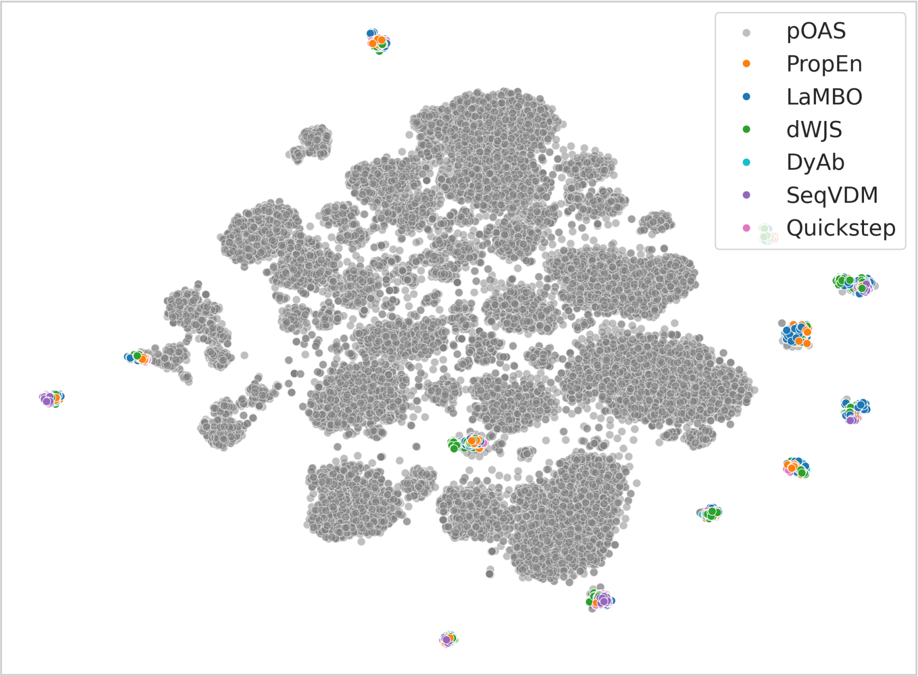


**Fig. B5**: t-SNE[84] plot of ESM-2 [54] embeddings of designs for all seeds plotted against paired OAS [57] (30k random sub-sample). Our seeds are mainly either on the fringes of this reference distribution or are completely OOD.

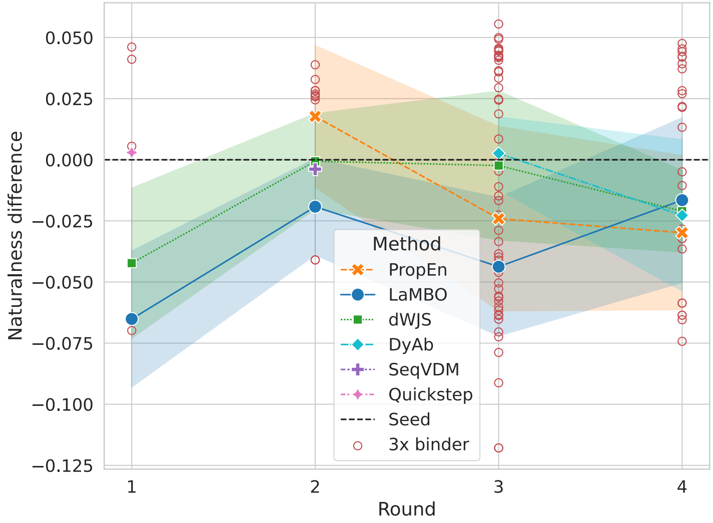


**Fig. B6**: Difference in naturalness score [85] (inverse perplexity of AntiBERTy PLM [19]) of the designs compared to “wild type” antibody seeds over the different rounds. Higher naturalness signifies that the language model deems the sequence to be likely, w.r.t. its training data. We can see that as number of 3x binders increases going from round 2 to 3 or 4 the naturalness generally decreases. This signifies that as the designs are improving they are drifting slightly away from a naive antibody distribution.

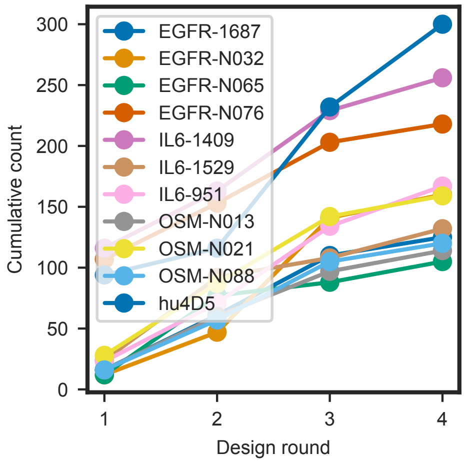


**Fig. B7**: Cumulative number of labeled (expression yield and binding affinity) designs in the neighborhood of each seed (<= 16 edit distance) over rounds

.

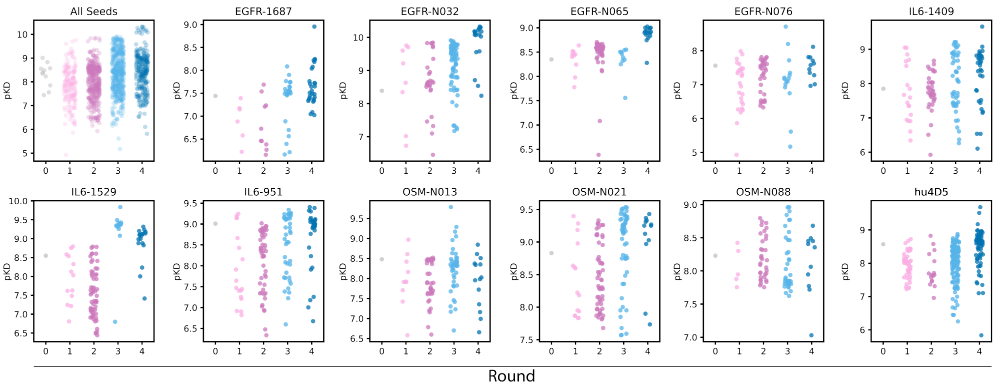


**Fig. B8**: Absolute $pK_D$ values for all designs, separated by seed. This is an alternative visualization of the design (blue) points in Figure 2d.

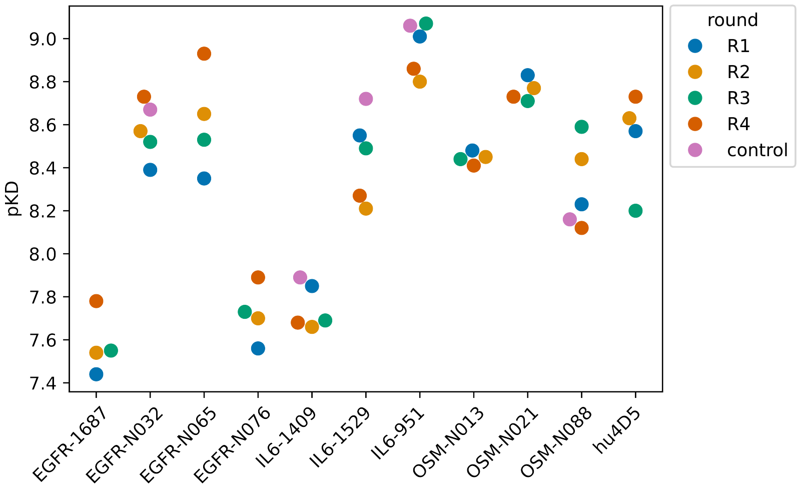


**Fig. B9**: Absolute $pK_D$ values of all seeds across all rounds to highlight reproducibility across biological replicates. This is an alternative visualization of the seed (pink) points in Figure 2d. Differences in reported $pK_D$ for each design are computed relative to the within-round seed value to normalize for batch effects. This is particularly important for the seed hu4D5-linear (see also Table A1), as its absolute affinity measured by antibody-capture SPR as described in Methods 4.4 can be highly variable.

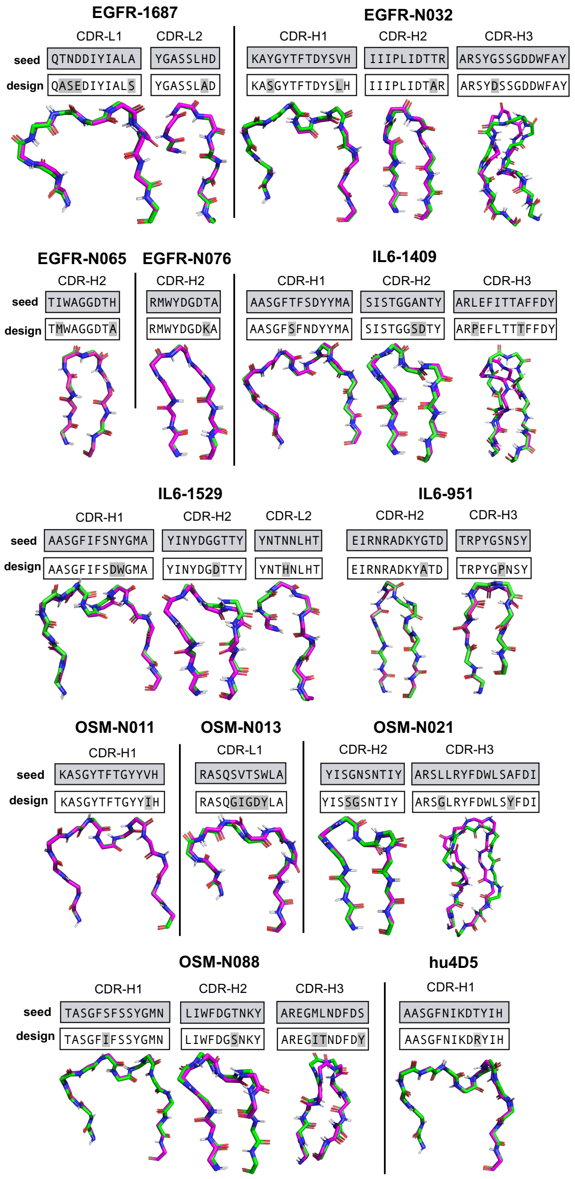


**Fig. B10**: CDR loop comparison between seeds and designs. Global alignment of seed (green) and affinity matured designs (magenta). All CDR loops (AHo definitions [46]) containing mutations between seed and design are shown (other mutations may be present in non-CDR regions).

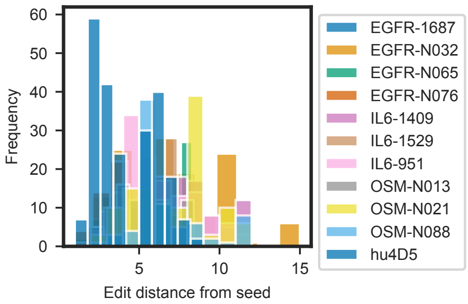


**Fig. B11**: Histogram of edit distances of all binding designs with respect to their respective seed sequences. Edit distances range from 1 to 14.

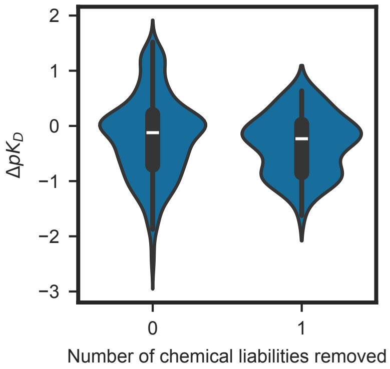


**Fig. B12**: Change in $pK_D$ versus number of chemical liabilities removed from a seed sequence.

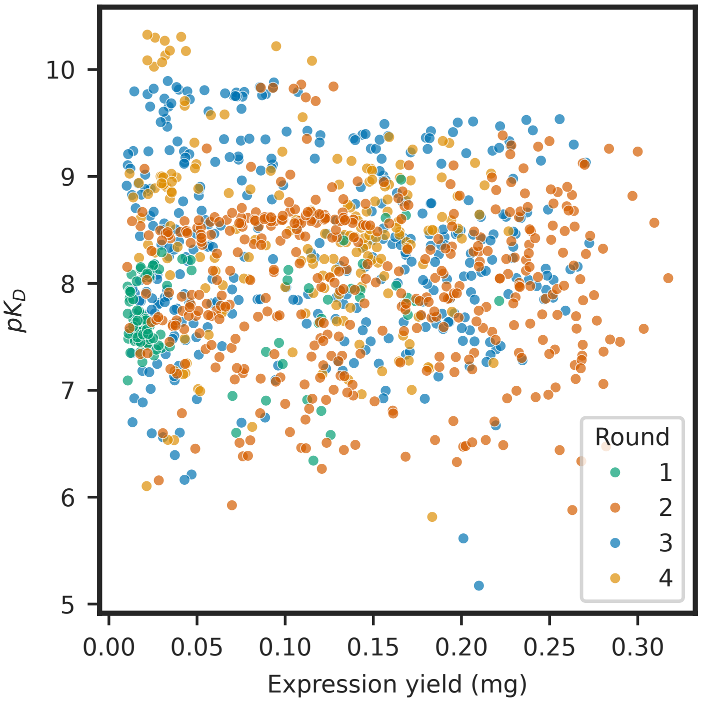


**Fig. B13**: Absolute expression yield and affinity values for all binders across all rounds, corresponding to the Pareto frontiers seen in Figure 3c. The expected variability for expression at 1 mL scale is approximately 10-15% based on historical data..

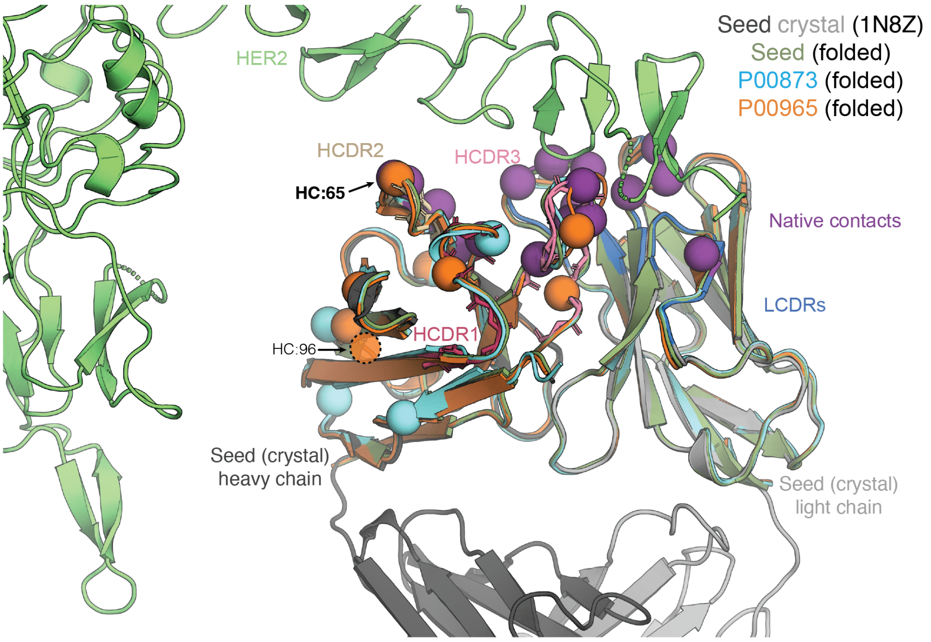


```
                              HCDR1                   HCDR2                                          HCDR3
EVQLVES-GGGLVQPGGSLRLSCAASGFNIKDTYIHWVRQAPGKGLEWVARIYPTNGYTRYADSVKGRFTISADTSKNTAYLQMNSLRAEDTAVYYCSRWGGDGFYAMDYWGQGTLVTVSS
EVQLVESDGGGLVQQGGSLRLSCAASGFNIKDRYIHWVRQAPGKGLEWVARIYPTNGYTRYADRVKGRFTISADTSKNTAYLQMFSLRAEDTAVYYCSRWGGDGFYAMDYWGQGTLVTVSS
EVQLVES-GGGLVQPGGSLRLSCAASGFNMKDTYIHWVRQAPGKGLEWVARIYPTQGYTRYADAVKGRFTISAPTSKNTAYLTMNS-RAEDTAVYYCSRWGGDGFYAFDYWGQGTLVTVSS
```

**Fig. B14**: Zoom-out of the trastuzumab (hu4D5)-HER2 complex crystal structure (PDBID: 1N8Z) with heavy/light chain colored in dark/light gray, HCDRs colored as in Fig 4, LCDRs in dark blue, native contacts in purple, with folded Fv structures (ABodyBuilder2) of the seed (dark green) and two designs (light blue and orange) aligned onto the crystal. A dashed transparent circle depicts the approximate location of the deletion in P00965. The folded structures exhibit very close similarities in all loops except for minor variability in HCDR3, indicating that the sequence changes are unlikely to have introduced dramatic structural rearrangements. Note that position 65 on the heavy chain (HC:65) is the only native seed contact that is directly mutated (in P00965 only). All other mutations are orthogonal to native contacts, with most localized to the framework region. The complete heavy-chain sequences for the seed and two designs are shown at bottom and color-coded by origin. There were no edits to the light chain in either of these two designs.

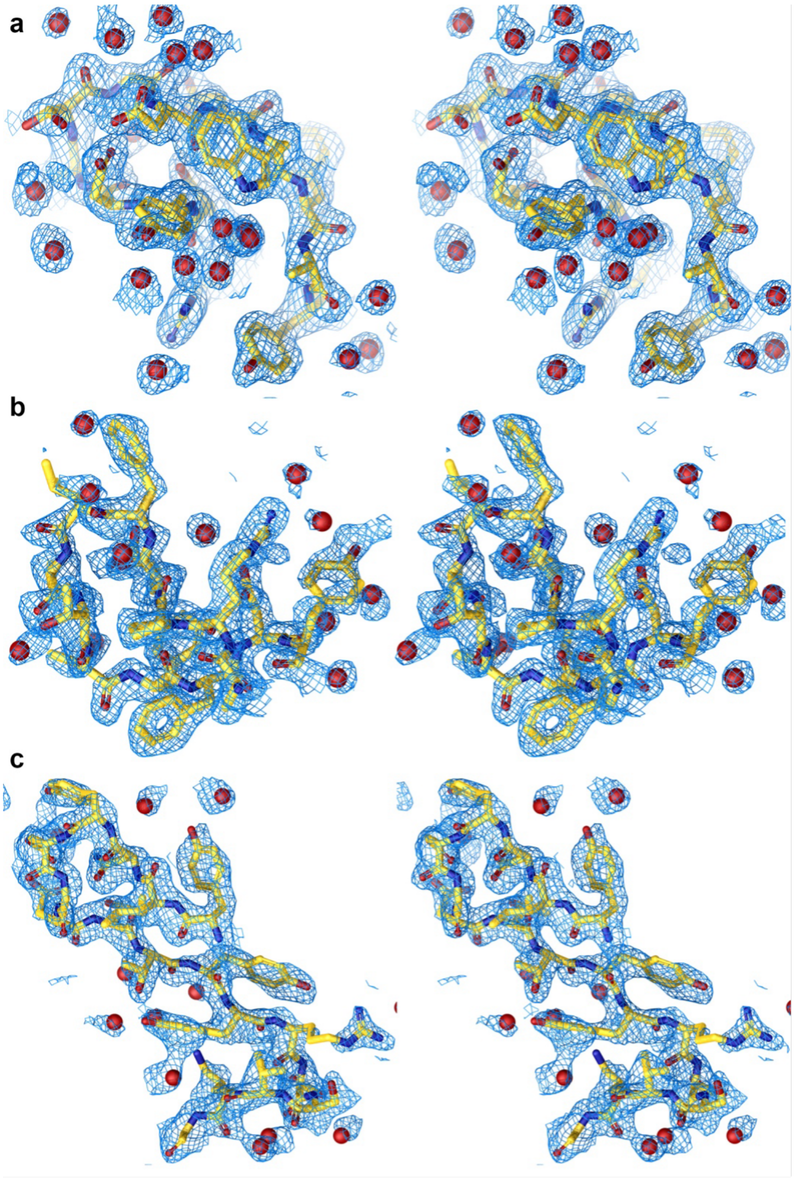


**Fig. B15**: Electron density maps. Divergent eye stereo images of the simulated annealing composite omit difference electron density maps, (2m$|F_o|$-D$|F_c|$) exp(i$\alpha$c), CDR H3 of Fabs **a** 9MVJ, **b** 9MUZ, **c** 9MUY (CDR H2), **d** 9MZC, **e** 9MZV, **f** 9N2V, **g** 9N7M, and **h** 9N7O, contoured at 1$\sigma$.

904

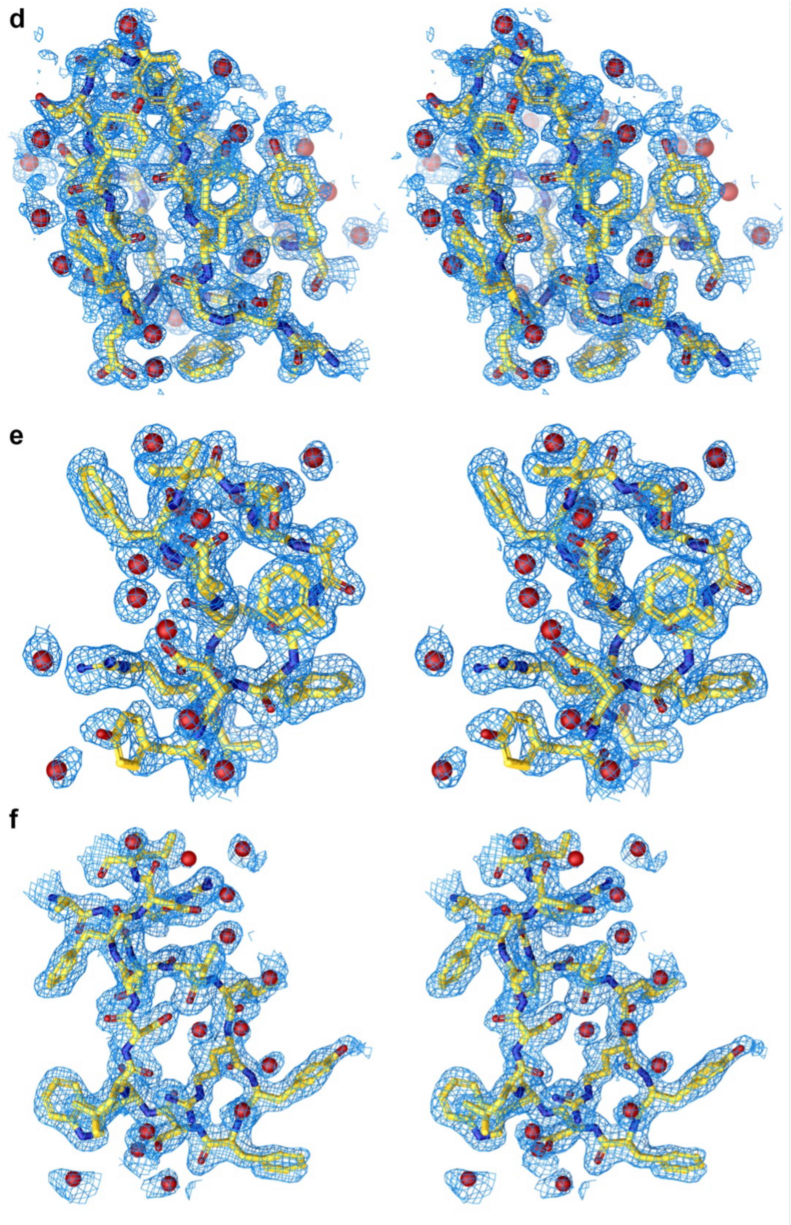

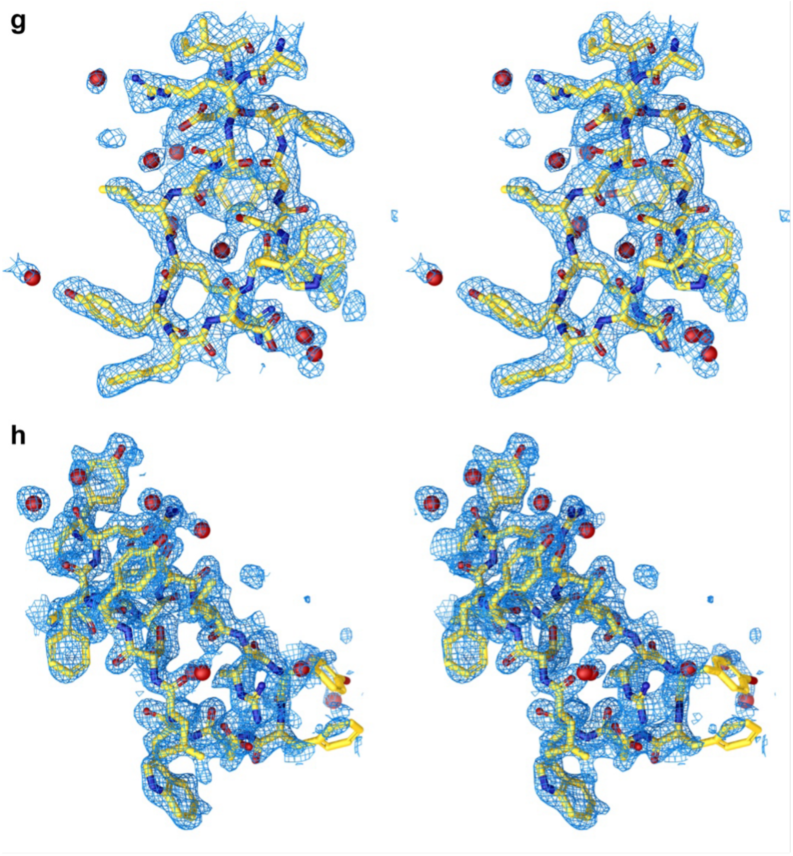
g
h

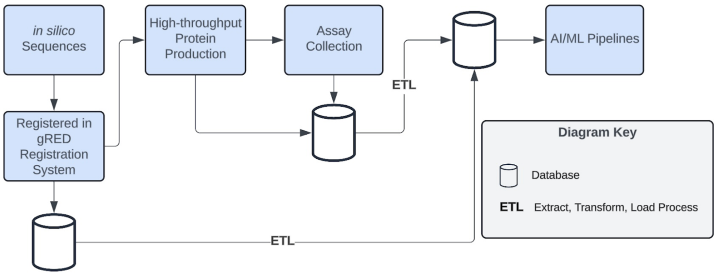


**Fig. B16**: Diagram of extract, transform, load process for data pipelines.
